# Nuclear acetyl-CoA production by acetyl-CoA synthetase enables dynamic histone acetylation and gene expression in *Plasmodium falciparum*

**DOI:** 10.64898/2026.07.16.738923

**Authors:** Reginald F. Akossi, Patty Chen, Jennifer J. Schwarz, Florent Dingli, Yoshiki Yamaryo-Botté, Catarina Rosa, Giulia Litta Modignani, Damarys Loew, Cyrille Y. Botté, Sebastian Baumgarten, Jessica M. Bryant

**Author notes:** Corresponding author, (JMB). Present address: Cell Biology of Host – Pathogen Interaction Laboratory, Gulbenkian Institute for Molecular Medicine, Oeiras, Portugal (Catarina Rosa). These authors contributed equally to this work (Reginald F. Akossi, Patty Chen).

## Abstract

A complex transcriptional cascade drives the two-day intraerythrocytic developmental cycle of the most virulent human malaria parasite, *Plasmodium falciparum*, in the human host. Genes are rapidly activated at specific times, then silenced as quickly. Transcriptional activity correlates with histone acetylation, which is modulated by acetyltransferases that use acetyl-CoA and deacetylases that produce acetate. Acetyl-CoA synthetase (ACAS) uses acetate to produce acetyl-CoA and has been implicated in histone acetylation, providing an interesting link between parasite metabolism and transcription. Here, we show that ACAS becomes enriched in the nucleus at a time during the life cycle when the parasite undergoes rapid growth and increased levels of transcription. We use mass spectrometry to show that ACAS inhibition results in a rapid and global depletion of histone acetylation, which leads to a general decrease in chromatin accessibility at gene promoters. These widespread alterations in chromatin composition disrupted the transcriptional cascade, resulting in cell cycle arrest. Our study provides evidence that ACAS plays an important nuclear role in the histone acetylation cycle and insight into the dynamic nature and essentiality of histone acetylation in the parasite’s complex transcriptional program driving infection of the human host.

## Introduction

Malaria caused by the *Plasmodium falciparum* parasite is a devastating disease afflicting sub-tropical regions (Weiss *et al*, 2025). The parasite intraerythrocytic developmental cycle (IDC) starts when a merozoite invades a human red blood cell where it is able to hide from the immune system while it replicates asexually (Grüring *et al*, 2011). The IDC is responsible for most clinical symptoms of malaria (Cowman *et al*, 2016) and is driven by a transcriptional cascade controlled via epigenetic mechanisms such as histone post translational modifications (Cortés & Deitsch, 2017; Painter *et al*, 2018). Histone acetylation is associated with open, transcriptionally permissive chromatin and is involved in gene activation (Cui *et al*, 2007; Rawat *et al*, 2020). Histone acetyltransferases (HATs) use acetyl-CoA to transfer an acetyl group to various histone residues, and histone deacetylases (HDACs) remove acetyl groups from histones, producing acetate. Thus, acetyl-CoA levels impact histone acetylation levels (Wellen *et al*, 2009).

Acetyl-CoA is involved in many other important metabolic processes including fatty acid synthesis and the TCA cycle (Pietrocola *et al*, 2015). In *P. falciparum*, acetyl-CoA can be produced by three different pathways. The first is via pyruvate dehydrogenase (PDH) localized to the apicoplast (aPDH) (Foth *et al*, 2005; Pei *et al*, 2010); however, as aPDH is dispensable during the IDC, this pathway is not critical to acetyl-CoA production in blood stage parasites (Pei *et al*, 2010; Cobbold *et al*, 2013; Swift *et al*, 2020). The mitochondrion is a major source of cellular acetyl-CoA via the branched-chain ketoacid dehydrogenase (BCKDH) complex, which acts as a mitochondrial PDH (mPDH) and was shown to play a role in the acetylation of histones and other proteins outside the mitochondrion (Cobbold *et al*, 2013; Oppenheim *et al*, 2014; Nair *et al*, 2023). Finally, acetyl-CoA synthetase (*Pf*ACAS) combines acetate (which can be scavenged from the host or generated in the cell) and coenzyme A to form acetyl-CoA. *Pf*ACAS is essential during the IDC, contributes significantly to the cytoplasmic and nuclear pools of acetyl-CoA (Cobbold *et al*, 2013; Schalkwijk *et al*, 2019), and can rescue parasite growth in lines deficient in mitochondrial acetyl-CoA production (Nair *et al*, 2023).

While it is clear that metabolites such as acetyl-CoA are needed in the nucleus for acetylation of histones and other proteins, a growing body of evidence supports specific and targeted nuclear roles for metabolic enzymes such as ACAS with regard to transcriptional regulation in mammals (Li *et al*, 2018). In mouse differentiating neurons, acetyl-CoA synthetase 2 (ACSS2) binds to active neuronal genes that are enriched in histone acetylation and whose transcription is down-regulated upon ACSS2 depletion (Mews *et al*, 2017). Moreover, a decrease in ACSS2 in the hippocampus of adult mice impairs long-term spatial memory (Mews *et al*, 2017). Another example is from human glioblastoma cells, where AMPK α phosphorylates ACSS2 at a C-terminal nuclear localization signal (NLS) in response to glucose deprivation, which leads to ACSS2 nuclear translocation (Li *et al*, 2017). In the nucleus, ACSS2 binds to a transcription factor and associates with promoters of genes involved in lysosome biogenesis and cell survival. There, ACSS2 generates local pools of acetyl-CoA that are then used to acetylate histones and enhance transcription of associated genes (Li *et al*, 2017). Thus, ACAS links changes in cellular metabolism to a transcriptional response in multiple cellular contexts.

As *P. falciparum* is reliant upon its host for nutrients but lacks canonical nutrient-sensing pathways (Serfontein *et al*, 2010; Babbitt *et al*, 2012; Miranda-Saavedra *et al*, 2012), *Pf*ACAS has emerged as an interesting candidate for epigenetic sensor. Indeed, *Pf*ACAS is found in the cytoplasm and nucleus, and its inhibition or knockdown led to a decrease in acetylation levels of certain histone residues; however, the stage at which *Pf*ACAS is nuclear and which histone residues are affected has been contested (Bryant *et al*, 2020; Prata *et al*, 2021; Summers *et al*, 2022; de Vries *et al*, 2022). We originally found *Pf*ACAS enriched in the chromatin of the upstream regions of *var* genes, which are part of a ∼60-member multigene family encoding an important variant surface antigen (Bryant *et al*, 2020). A more recent study by Prata *et al*. performed chromatin immunoprecipitation and sequencing (ChIP-seq) and found peaks of *Pf*ACAS associated with over 3,000 genes that are expressed at all stages of the life cycle (Prata *et al*, 2021). The same study found that knockdown or inhibition of *Pf*ACAS led to a decrease in active *var* gene transcription (Prata *et al*, 2021). While these two studies show a direct interaction of *Pf*ACAS with chromatin, it is unclear whether this metabolic enzyme plays a direct transcriptional role at specific genes or contributes to transcriptional activity in a general manner via global histone acetylation.

To investigate the nuclear role of *Pf*ACAS, we undertook an omics-based comprehensive functional characterization using a parasite-specific inhibitor first described in (Summers *et al*, 2022). We found that *Pf*ACAS inhibition led to a global decrease in histone acetylation and chromatin accessibility accompanied by disruption of the IDC transcriptional cascade, indicating that parasite chromatin composition and transcription are intimately linked to nutrient availability. Moreover, manipulation of *Pf*ACAS revealed a highly dynamic nature of histone acetylation. While our data suggest a general nuclear role for *Pf*ACAS in providing acetyl-CoA for histone acetylation and transcriptional activation, we also observed a very specific enrichment of *Pf*ACAS in heterochromatic, subtelomeric regions that are important for genome architecture and contain genes that were up-regulated upon *Pf*ACAS inhibition, hinting at a non-canonical role.

## Results

### Nuclear localization of PfACAS is highest in trophozoites and is responsive to glucose levels

To study the nuclear role of *Pf*ACAS, we first set out to determine at which stage(s) of the IDC *Pf*ACAS is most enriched in the nucleus. We generated a parasite strain in which endogenous *Pf*ACAS is fused to a hemagglutinin (HA) tag (**Fig. EV1A**). A western blot time course revealed that while *Pf*ACAS is present at constant levels in the cytoplasm over the course of the IDC, it is most enriched in the nucleus in trophozoite and schizonts stages (**Fig. 1A)**. Localization of *Pf*ACAS in both the cytoplasm and nucleus in trophozoites was further confirmed by immunofluorescence (**Fig. 1B**) and a separate strain in which *Pf*ACAS is tagged with GFP (**Fig. EV1A,B**).

**Figure 1:**
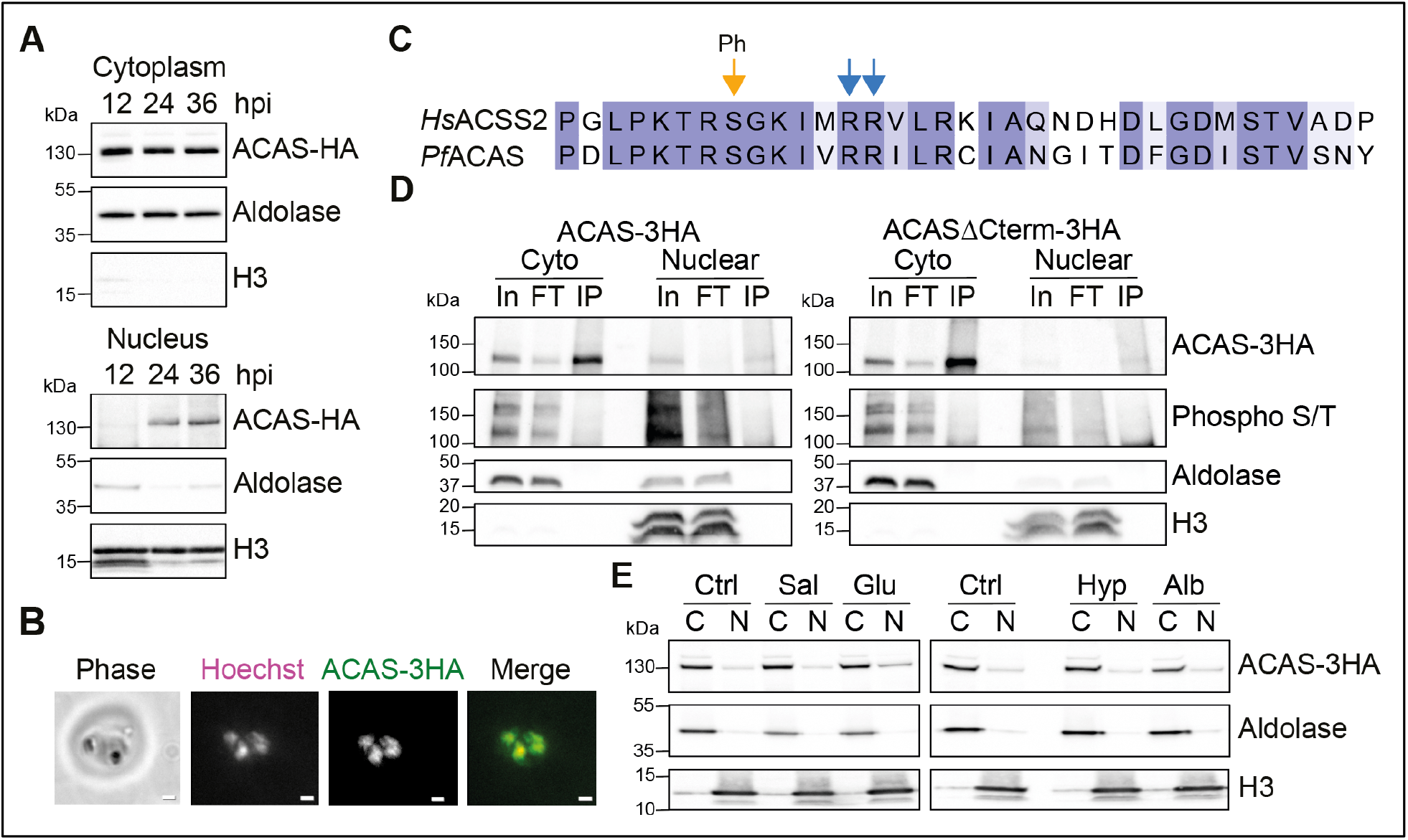
Nuclear localization of *Pf*ACAS increases in trophozoites and is responsive to glucose levels **A.** Western blot analysis of nuclear and cytoplasmic fractions from a synchronous population of *Pf*ACAS-3HA (detected with an anti-HA antibody) parasites over the course of the IDC (hours post invasion indicated at top). Antibodies against aldolase and histone H3 are used as cytoplasmic and nuclear controls, respectively. Molecular weights are shown to the left. **B.** Immunofluorescence assay of fixed RBCs infected with trophozoite *Pf*ACAS-3HA parasites. DNA was stained with Hoechst (magenta), and *Pf*ACAS-3HA was detected with anti-HA (green) antibody. Scale bars equal 2 μm. **C.** Alignment of the C-termini of human ACSS2 and *Pf*ACAS. Intensity of purple indicates level of conservation. The serine residue (amino acid 946 in *Pf*ACAS) phosphorylated by AMPK and the two arginine residues required for nuclear translocation of ACSS2 are indicated with orange and blue arrow heads, respectively (Li *et al*, 2017). **D.** Western blot analysis with an anti-HA or anti-phosphorylated serine/threonine antibody of cytoplasmic or nuclear extracts from synchronous ACAS-3HA or ACASΔCterm-3HA trophozoites (24hpi) before (input, “In”) and after (flow-through, “FT”) immunoprecipitation (IP) with an antibody against HA. Antibodies against aldolase and histone H3 are used as a cytoplasmic and nuclear controls, respectively. Molecular weights are shown to the left. **E.** Western blot analysis of nuclear (N) and cytoplasmic (C) fractions from a synchronous population of trophozoite *Pf*ACAS-3HA (detected with an anti-HA antibody) parasites cultured for one hour with DMSO (Ctrl), salicylate (Sal), or media without glucose (Glu), hypoxanthine (Hyp), or albumax (Alb). Antibodies against aldolase and histone H3 are used as cytoplasmic and nuclear controls, respectively. Molecular weights are shown to the left.

Since our data suggested shuttling of *Pf*ACAS between the cytoplasm and nucleus, we set out to identify factors that could facilitate *Pf*ACAS nuclear translocation. We performed *Pf*ACAS immunoprecipitation and mass spectrometry (IP-MS) using cytoplasmic and nuclear fractions from trophozoites (**Supp. Data 1,2**). One of the most highly enriched proteins in the *Pf*ACAS IPs (versus wildtype control) was *Pf*ACAS, validating our strain and method (**Fig. EV1C,D, Supp. Data 1,2**). Gene Ontology (GO) analysis of proteins that were significantly abundant in the *Pf*ACAS IPs revealed an enrichment of terms relating to glycolysis (**Supp. Data 3**). In fact, most enzymes in the glycolytic pathway were enriched, which could indicate the physical association of two different acetyl-CoA-producing pathways (**Fig. EV1C**). Terms related to RNA Polymerase II transcription, which include subunits of RNA Pol II and the FACT complex and transcription elongation factors, were also enriched in the *Pf*ACAS IP (**Fig. EV1D, Supp. Data 1-3**), suggesting association with active transcriptional machinery in the nucleus. Indeed, the enrichment of several other chromatin-associated proteins such as Bromodomain Protein 1, ApiAP2 DNA-binding factors, the SET6 histone methyltransferase, and putative remodelers ISWI and microrchidia (MORC) hint at the presence of *Pf*ACAS in specific transcriptional complexes (**Fig. EV1D, Supp. Data 2**). In addition, several importins, exportins, and nuclear pore complex subunits were found to be enriched in the *Pf*ACAS IPs, suggesting targeted nuclear import and export of *Pf*ACAS (**Fig. EV1C,D, Supp. Data 1,2**).

Next, we investigated whether *Pf*ACAS nuclear translocation could be controlled via AMPK phosphorylation, similarly to ACSS2 in human cancer cells (Li *et al*, 2017). *P. falciparum* does not have a canonical AMPK heterotrimeric complex, but it encodes a serine/threonine kinase, KIN, with limited homology to mammalian AMPKα (Mancio-Silva *et al*, 2017). *Pf*KIN is involved in sensing changes in host nutrient availability and triggering the parasite’s transcriptional response (Mancio-Silva *et al*, 2017). Interestingly, *Pf*ACAS has a C-terminal nuclear localization signal that is very similar to the one found in human ACSS2 that is phosphorylated by AMPK (**Fig. 1C**). Multiple attempts to mutate this putative phosphorylation site at the endogenous locus were unsuccessful, suggesting that these residues are important for the function of *Pf*ACAS. However, when we expressed *Pf*ACASΔCterm-3HA that was mutated at this putative phosphorylation site (S946A_R951A_R952A) from an episome, its nuclear localization was not affected in trophozoites (**Fig. 1D**). Moreover, we did not observe significant phosphorylation of wildtype or *Pf*ACASΔCterm in the cytoplasm or the nucleus of trophozoites (**Fig. 1D**). Curiously, we were never able to express wildtype *Pf*ACAS from an episome, suggesting that *Pf*ACAS overexpression is lethal and that the mutated *Pf*ACAS is not functional.

While phosphorylation of *Pf*ACAS, at least to the extent that is detectable by western blot, does not seem to play a role in its nuclear translocation, it is possible that other pathways could be involved in cytoplasmic-nuclear shuttling. To determine if *Pf*ACAS nuclear translocation is affected by changes in nutrient availability, we cultured parasites for one hour in media without glucose, without hypoxanthine (essential for nucleic acid metabolism (Desai, 2013)), without albumax (source of various important lipids (Srivastava *et al*, 2007)), or with salicylate (an AMPK activator used previously to manipulate *Pf*KIN (Mancio-Silva *et al*, 2017)). Only glucose withdrawal led to a slight increase in nuclear *Pf*ACAS levels, suggesting that *Pf*ACAS shuttling could be responsive to host glucose levels (**Fig. 1E**).

### PfACAS associates with heterochromatinized regions and proteins involved in subtelomeric structure

Because *Pf*ACAS becomes enriched in the nuclei of trophozoites, we performed ChIP-seq in this stage of the IDC in two separate strains: *Pf*ACAS-3HA and *Pf*ACAS-GFP (**Fig. EV1A)**. In general, *Pf*ACAS was enriched at low levels across the genome in what appeared to be non-specific, potentially background-level binding (**Fig. 2A**). Curiously, the most significant consensus peaks were found upstream of *upsB var* genes in subtelomeric regions of the genome (**Fig. 2A-C, Supp. Data 4**). These *Pf*ACAS binding sites overlap with or are in close proximity to those of several other factors including AP2-P, MORC, SIP2, and TRZ at a region we recently found to be important for subtelomeric chromatin structure and *var* gene clustering (Flueck *et al*, 2010; Bertschi *et al*, 2017; Singh *et al*, 2025) (**Fig. 2B**). Indeed, interaction between *Pf*ACAS and these other factors is supported by 1) our previous proteomic characterization of *var* gene chromatin (Bryant *et al*, 2020) and 2) our IP-MS analysis of *Pf*ACAS as well as an IP-MS analysis of MORC (Singh *et al*, 2024) (**Supp. Data 2**). These subtelomeric regions are heterochromatinized via heterochromatin protein 1 (HP1) and are generally devoid of histone acetylation, suggesting that there might be elevated levels of acetylation of a non-histone protein or that *Pf*ACAS plays a divergent role at these particular subtelomeric sites (**Fig. 2B,C**).

**Figure 2:**
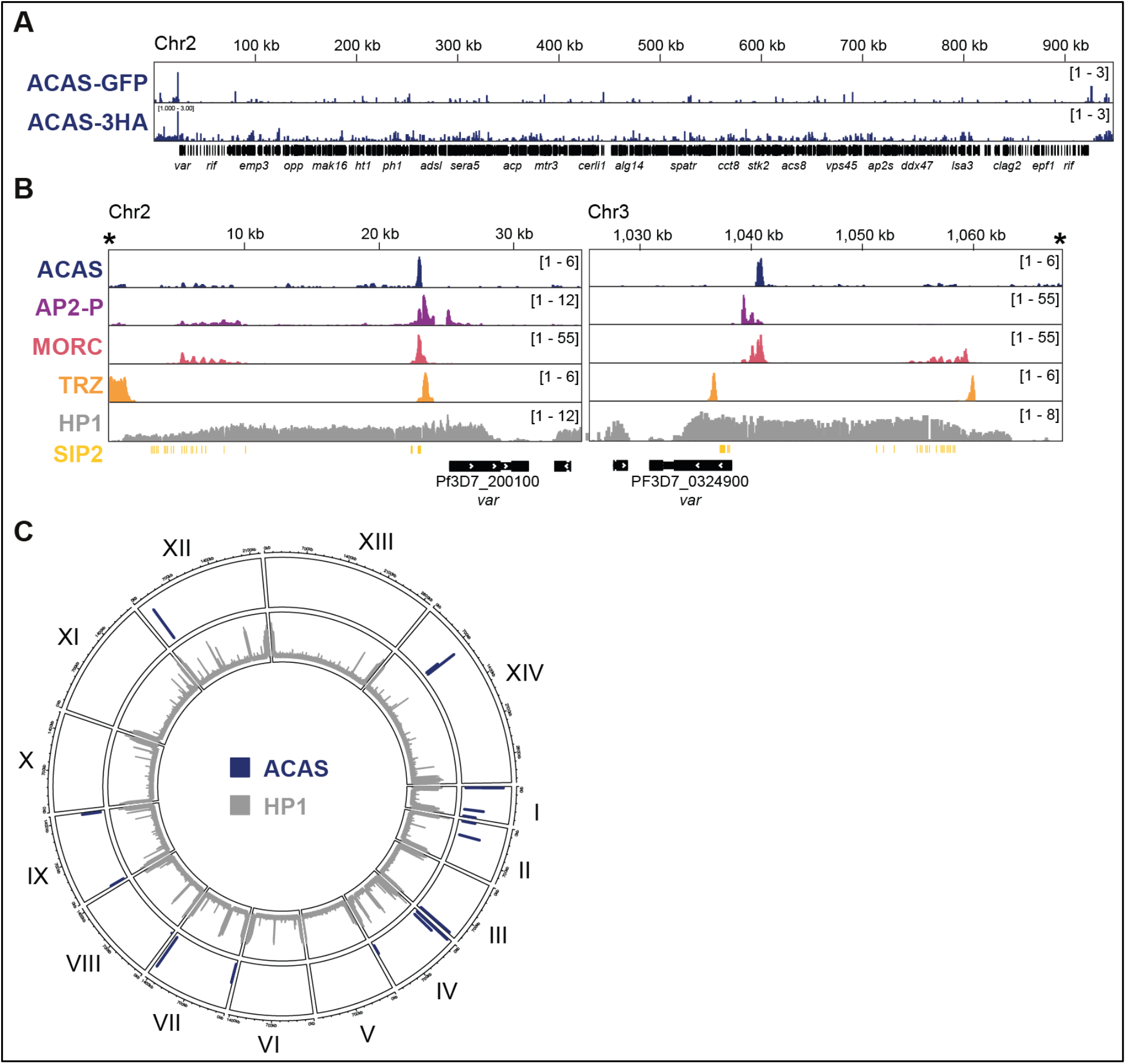
*Pf*ACAS is enriched in subtelomeric regions upstream of *var* genes **A.** ChIP-seq data (ChIP/Input) of *Pf*ACAS-GFP and *Pf*ACAS-3HA on chromosome 2. The x-axis is DNA sequence, with genes represented with black boxes. **B.** ChIP-seq data (ChIP/Input) of ACAS in trophozoites and AP2-P (Singh *et al*, 2025), MORC (Singh *et al*, 2025), TRZ (Bertschi *et al*, 2017), and HP1 (Carrington *et al*, 2021) in schizonts at the subtelomeric regions on the left arm of chromosome 2 and the right arm of chromosome 3. Putative SIP2 binding sites (SPE2 sequence) (Flueck *et al*, 2010) are indicated with vertical yellow lines. The x-axis is DNA sequence, with the *upsB var* gene represented by a black box with white arrowheads to indicate transcription direction. The telomere is indicated with an asterisk. **C.** Circos plot of consensus ChIP-seq peaks of *Pf*ACAS in trophozoites (blue) (**Supp. Data 4**). The 14 chromosomes are represented circularly by the outer bars, with chromosome number indicated in roman numerals and chromosome distances (Mbp) indicated in Arabic numerals. HP1 enrichment ratio [ChIP/input, (Carrington *et al*, 2021)] is shown as average reads per million (RPM) over bins of 1,000 nucleotides. The maximum y-axis value is 11.3083.

### PfACAS plays a role in global histone acetylation

As we observed *Pf*ACAS enriched in the nucleus and at specific loci in the genome, we wanted to determine its role in chromatin composition and specifically histone acetylation. Previous studies using a human *Hs*ACSS2 inhibitor (Prata *et al*, 2021) or a Plasmodium-specific ACAS inhibitor (Summers *et al*, 2022) demonstrated impaired parasite growth and a decrease in histone acetylation levels; however, only a few histone residues were assayed with western blot analysis. Thus, we set out to determine if *Pf*ACAS inhibition affects a specific histone residue or global histone acetylation. First, we obtained the same parasite-specific inhibitor – MMV019721 – used by (Summers *et al*, 2022) and confirmed that treatment of synchronized wildtype trophozoite cultures for three hours with inhibitor (2.5µM) did not immediately lead to parasite death but resulted in a decrease in levels of histone acetylation with western blot analysis using antibodies against H3K9ac, H3 pan acetylation, and H4 pan acetylation (**Fig. 3A**).

**Figure 3:**
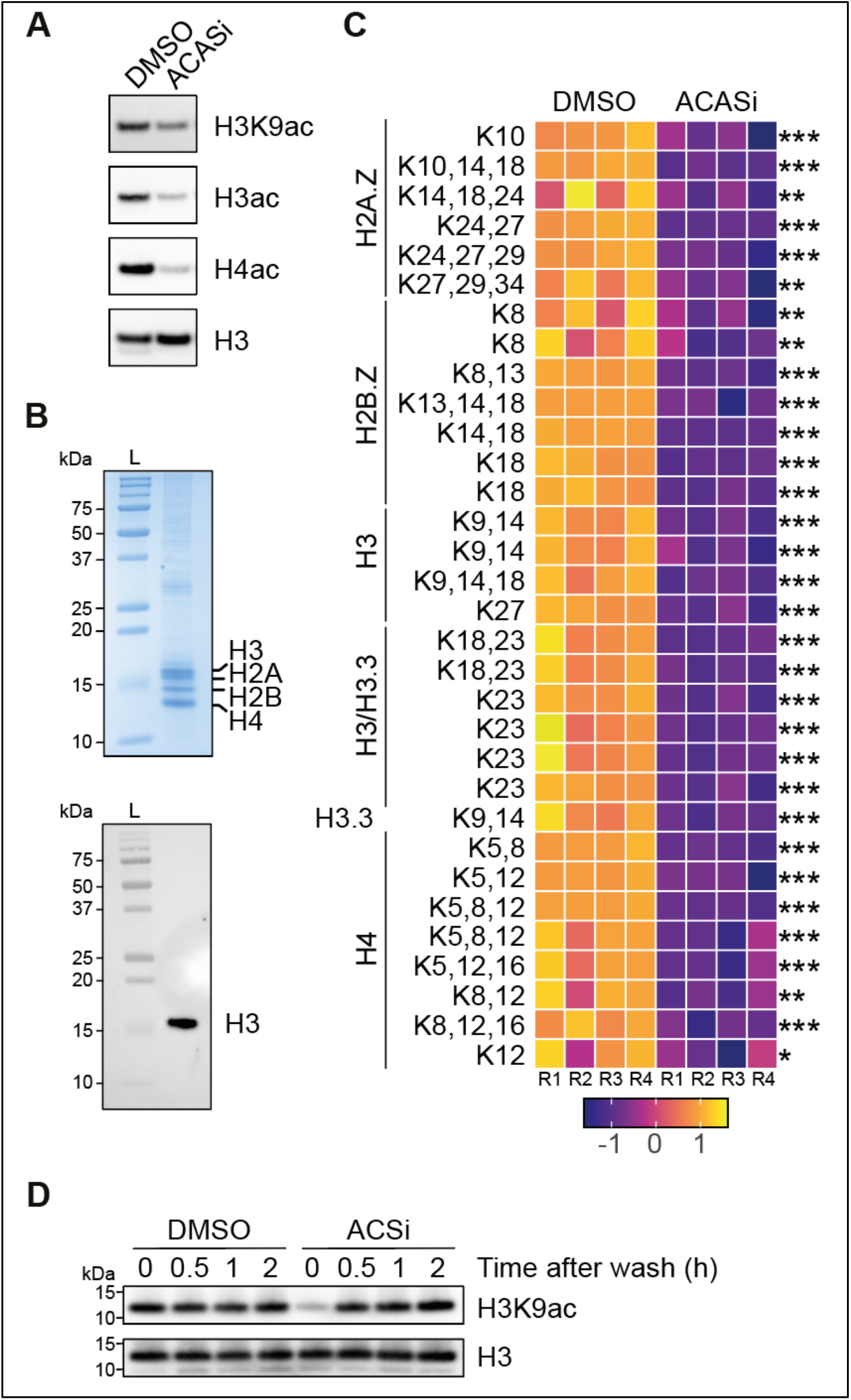
ACAS inhibition induces global reduction in histone acetylation. **A.** Western blot analysis of nuclear extracts from synchronous wildtype trophozoites (21 hpi) treated for three hours with DMSO or ACASi at 2.5µM. Histone H3, H3K9ac, histone H3 pan-acetyl (H3ac), and histone H4 pan-acetyl (H4ac) antibodies were used. **B.** Protein gel (upper panel) and western blot analysis using a histone H3 antibody (lower panel) of histone extracts from synchronous wildtype trophozoites (24 hpi). Molecular weights from the ladder (L) are shown to the left. Protein bands corresponding to histones H3, H2A, H2B, and H4 are indicated on the protein gel. **C.** Acetylation of peptides at indicated lysine residues of histones H2A.Z, H2B.Z, H3, H3.3, and H4 in four replicates each of DMSO-versus ACASi-treated synchronized wildtype trophozoites (24 hpi) obtained by mass spectrometry. The color scale represents the z-score of the log_2_ - transformed ratio of acetylated peptide abundance (MS1 total area) relative to its corresponding normalized histone protein abundance. Some lysine acetylation sites are represented multiple times, as several distinct peptides covered this specific site. Student t-test (*** p ≤ 0.001, ** p ≤ 0.01, * p ≤ 0.05, ns > 0.05). See**Supp. Data 5**. **D.** Western blot analysis of nuclear extracts from synchronous wildtype trophozoites that were treated for three hours with DMSO or ACASi (“0h”), then washed and sampled at different times after the wash. Histone H3 and H3K9ac antibodies were used. Molecular weights are shown to the left.

To have a global view of how *Pf*ACAS inhibition affects histone acetylation, we repeated the ACASi treatment and performed mass spectrometry on purified histones (**Fig. 3B**), focusing the analysis on histone acetylation (**Supp. Data 5**). While we obtained more than 87% coverage of all histone proteins, we were not able to distinguish between H3 and H3.3 for certain peptides due to the enzymatic digestion pattern and homology. Interestingly, we found acetylated peptides from H2A.Z and H2B.Z, but not from H2A and H2B (**Fig. 3C, Supp. Data 5**). This trend was also observed by (Trelle *et al*, 2009) and supports studies that found nucleosomes containing these histone variants to be enriched in the promoter regions of genes where histone acetylation is also enriched (Bártfai *et al*, 2010; Hoeijmakers *et al*, 2013; Petter *et al*, 2013).

Strikingly, histone peptides containing acetylation at every histone residue analyzed were less abundant in the ACASi-treated parasites compared to the DMSO-treated parasites (**Fig. 3C, Supp. Data 5**). These data show that ACAS inhibition, even for as little as three hours, leads to a global and significant decrease in histone acetylation. To test the dynamic nature of histone acetylation, we treated synchronized trophozoite wildtype parasites with ACASi for three hours, washed off the drug, and performed a western blot time course analysis to determine how quickly histone acetylation levels returned to normal. After only 30 minutes, histone H3K9ac levels returned to normal (**Fig. 3D**). Taken together, these findings suggest that *Pf*ACAS is essential for most, if not all, histone acetylation post-translational modifications (PTM) and that histone acetylation itself is extremely dynamic.

We wanted to compare the nuclear and extra-nuclear effects of *Pf*ACAS inhibition. As ACAS is known to play a role in lipid synthesis and elongation in a related apicomplexan parasite, *Toxoplasma gondii* (Dubois *et al*, 2018), we again treated synchronized wildtype trophozoites with ACASi for three hours and performed lipid mass spectrometry (**Supp. Data 6**). Our analysis revealed significant reductions in the abundance of only a few types of lipids, including di-and triacylglycerols and phosphatidylglycerol, which supports a dominant role for aPDH in generating acetyl-CoA for *de novo* lipid synthesis (**Fig. EV2A**). A higher (but not significant) proportion of 16C and 18C monoacyl glycerol (MAG) molecules relative to other MAGs in ACASi-treated parasites suggests a minor role for *Pf*ACAS in lipid elongation in the endoplasmic reticulum (**Fig. EV2B**). However, these changes were not as dramatic as the effects seen on histone acetylation, suggesting that mPDH compensates for loss of *Pf*ACAS activity in lipid elongation.

### PfACAS inhibition freezes the transcriptional cascade of the IDC

Because ACAS inhibition leads to massive and rapid depletion of histone acetylation in a way that might not be possible by inhibiting individual or even certain classes of histone acetyltransferases, it is a useful tool for studying the role of histone acetylation in chromatin accessibility and transcription. To determine the effect of *Pf*ACAS inhibition (and subsequent massive depletion of histone acetylation) on parasite transcription, we performed RNA sequencing in synchronous wildtype trophozoites that were treated with DMSO or ACASi for three hours. Despite the depletion of histone acetylation, there were approximately the same number of genes whose transcripts increased (927) or decreased (1,123) significantly in abundance by more than two-fold upon *Pf*ACAS inhibition relative to DMSO treatment (**Fig. 4A, Supp. Data 7**). The genes with increased transcript abundance were significantly enriched for GO terms relating to antigenic variation (**Supp. Data 8**), as almost all *var* and *rifin* transcripts, whose genes are normally heterochromatinized, were more abundant (**Fig. 4A, Supp. Data 7**). The genes with decreased transcript abundance were significantly enriched for GO terms relating to DNA repair and replication, as well as several biosynthetic and metabolic processes (**Supp. Data 8**).

**Figure 4:**
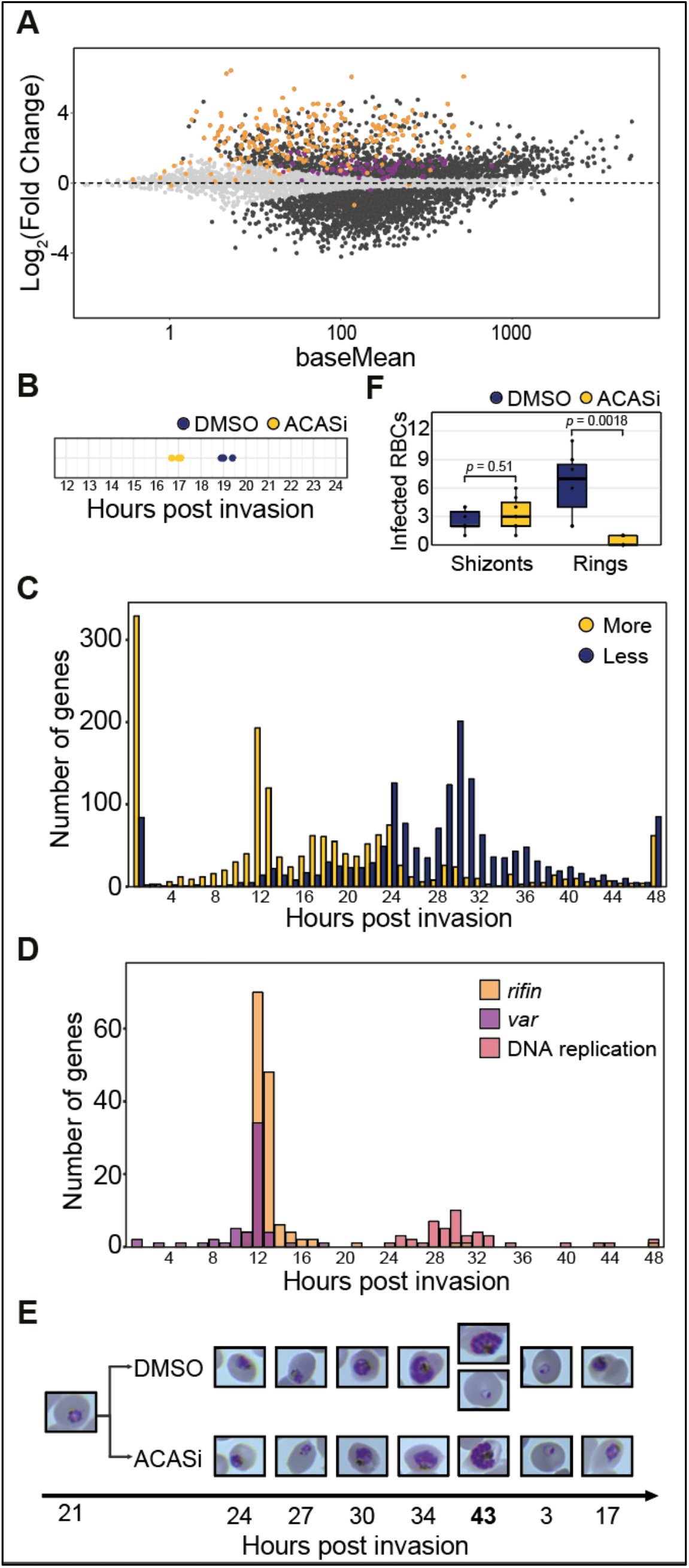
PfACAS inhibition causes a reversible stop in parasite transcriptional program and cycle progression. **A.** Differential expression analysis of ACASi-treated wildtype trophozoites (See **Supp. Data 7**). MA plot of log_2_(ACASi/DMSO) plotted over the mean abundance of each gene. Transcripts that were significantly higher (above x axis) or lower (below x axis) in abundance after treatment are highlighted in dark grey (q ≤ 0.05). Transcripts that showed no significant change are light grey, *rifin* genes are orange, and *var* genes are purple. Three technical replicates were used for DMSO-and ACASi-treated parasites. P values were calculated with a Wald test for significance of coefficients in a negative binomial generalized linear model as implemented in DESeq2 (Love *et al*, 2014). q = Benjamini–Hochberg adjusted P value. **B.** Cell cycle progression (hours post invasion on x-axis) estimation of wildtype trophozoites treated with DMSO (blue) or ACASi (yellow) for three hours. RNA-seq data from synchronized parasites harvested at 24 hpi were compared to microarray data from (Bozdech *et al*, 2003) as in (Lemieux *et al*, 2009) to determine the approximate time point in the IDC. Replicates are represented with circles. **C.** Frequency plot showing the time in the red blood cell cycle (hours post invasion of the red blood cell) of peak transcript level [comparison to transcriptomics time course in (Painter *et al*, 2018)] for gene transcripts that are significantly two-fold less (blue) or more (yellow) abundant following ACASi treatment in trophozoites. **D.** Frequency plot showing the time in the red blood cell cycle (hours post invasion of the red blood cell) of peak transcript level [comparison to transcriptomics time course in (Painter *et al*, 2018)] for *var* (purple) or *rifin* (orange) genes or genes whose products are involved in DNA replication (pink). **E.** Giemsa-stained blood smears with wildtype parasites before (21 hpi) and after (24 hpi and on) a three-hour treatment with DMSO or ACASi. **F.** Quantification of red blood cells infected with schizonts or rings per field of view at 43 hpi from Fig. 4E. Each field of view is represented with a circle. Boxes represent the median and IQR, and whiskers represent maximum and minimum values IQR. P values from Wilcoxon test are indicated.

We found no obvious functional pattern amongst the thousands of gene transcripts that were more or less abundant upon ACAS inhibition. To determine if these large differences in transcription were due to a difference in cell cycle progression between the DMSO-and ACASi-treated trophozoites, we calculated the transcriptional age of each condition. We found that the ACASi-treated parasites were 2-3 hours behind in the IDC compared to the DMSO-treated parasites (**Fig. 4B**). Indeed, genes that were significantly upregulated more than two-fold upon ACAS inhibition tend to be expressed most highly earlier in the IDC during the transition from rings to trophozoites, and genes that were significantly downregulated more than two-fold tend to be expressed most highly later in the IDC during the transition from trophozoites to schizonts (**Fig. 4C**). A delay in cell cycle progression at least partially explains the decreased abundance of transcripts encoding proteins involved in DNA replication and lipid-related metabolic and biosynthetic processes, as expression of the corresponding genes increases from trophozoites to schizonts (Painter *et al*, 2018) (**Fig. 4D**). The opposite is then true for the increased abundance of *var* and *rifin* transcripts, as these genes are expressed earlier in ring stages and then become repressed in trophozoites (Scherf *et al*, 1998; Kyes *et al*, 1999) (**Fig. 4D**).

To investigate correlative morphological differences in cell cycle progression, we treated parasites for three hours with ACASi, washed the drug off, and monitored with Giemsa staining for another cycle. While it was difficult to discern such a small (2-3 hours) difference in cell cycle progression at most stages, ACASi-treated parasites showed delayed schizont bursting and red blood cell reinvasion compared to DMSO-treated parasites (**Fig. 4E,F**). These data suggest that ACASi treatment freezes parasites transcriptionally during the IDC, which is reversible at least after short-term treatment.

### PfACAS inhibition and depletion of histone acetylation lead to a decrease in chromatin accessibility

Histone acetylation is believed to inherently loosen the attraction between the positively charged histones and negatively charged DNA, leading to less compact, more accessible chromatin (Cui *et al*, 2007). In *P. falciparum*, the Tn5 transposase used to perform the Assay for Transposase-Accessible Chromatin followed by sequencing (ATAC-seq) preferentially targeted close to the transcription start sites (TSS) of active genes that were also enriched in H3K9ac (Ruiz *et al*, 2018; Toenhake *et al*, 2018). To determine whether ACAS inhibition and depletion of histone acetylation changed chromatin accessibility, we treated synchronous trophozoites with ACASi as above and performed ATAC-seq. The peaks of ATAC-seq enrichment in our control (DMSO-treated) replicates showed significant overlap and concordance with ATAC-seq peaks at a similar time point from a different study, validating our methodology and analysis (Toenhake *et al*, 2018) (**Fig. 5A-C, Supp. Data 9**).

**Figure 5:**
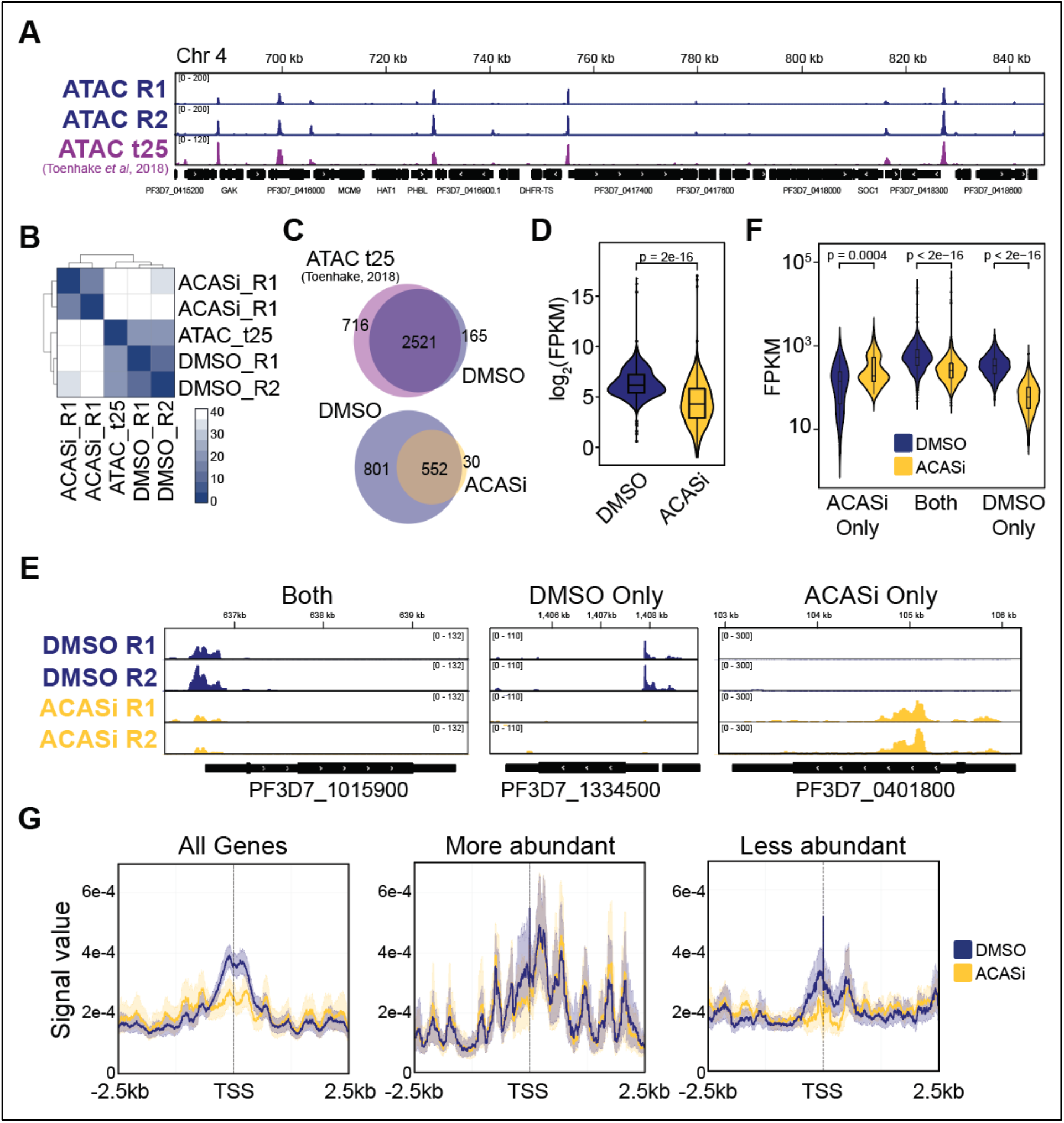
PfACAS inhibition causes a general loss of chromatin accessibility. **A.** ATAC-seq data from two control replicates (R1 and R2) of synchronized trophozoites (24 hpi) from this study and ATAC-seq data from a closely corresponding stage (t25) from (Toenhake *et al*, 2018) show chromatin accessibility [*y*-axis = ATAC-seq (RPM)/gDNA (RPM)]. The *x*-axis is DNA sequence from chromosome 4, with genes represented by black boxes with white arrowheads to indicate transcription direction. **B.** Comparison (Euclidean distance) of data from replicates of DMSO-and ACASi-treated trophozoites from this study and those from (Toenhake *et al*, 2018). See **Supp. Data 9**. **C.** Venn diagram showing overlap between consensus ATAC-seq peaks from two replicates of DMSO-or ACASi-treated synchronized trophozoites (24 hpi) from this study and ATAC-seq peaks from a closely corresponding stage (t25) from (Toenhake *et al*, 2018). **D.** Average (from two replicates) peak enrichment in DMSO-and ACASi-treated synchronized trophozoites (24 hpi). Boxes represent the median and IQR, and whiskers represent ±1.5× IQR. P value from a Wilcoxon rank sum test with continuity correction is indicated. **E.** Representative examples of genes with an associated ATAC-seq peak in both DMSO-and ACASi-treated, only in DMSO-treated, or only in ACASi-treated trophozoites. ATAC-seq data from two control replicates (R1 and R2) of DMSO-and ACASi-treated synchronized trophozoites (24 hpi) show chromatin accessibility [*y*-axis = ATAC-seq (RPM)/gDNA (RPM)]. The *x*-axis is DNA sequence, with genes represented by black boxes with white arrowheads to indicate transcription direction. **F.** Average (from two replicates) transcript abundance upon DMSO (blue) or ACASi (yellow) treatment of genes with an associated peak in both DMSO-and ACASi-treated, only in DMSO-treated, or only in ACASi-treated trophozoites. Boxes represent the median and IQR, and whiskers represent ±1.5× IQR. P value from a Wilcoxon rank sum test with continuity correction is indicated. **G.** Metagene analysis of average (two replicates) ATAC-seq signal in DMSO-(blue) or ACASi-treated (yellow) trophozoites from 2.5 kb upstream to 2.5 kb downstream of the transcription start site (TSS) for all genes, as well as genes whose transcripts are significantly two-fold more or less abundant after ACAS inhibition.

There were fewer consensus peaks in the ACASi-treated parasites (582 peaks) than in the DMSO-treated parasites (1,353 peaks) (**Fig. 5C**, **Supp. Data 9**). While there were 801 consensus peaks that were only found in DMSO-treated parasites, there were only 30 found only in ACASi-treated parasites (**Fig. 5C**). The average ATAC-seq peak enrichment in the DMSO-treated parasites was significantly higher than that in the ACASi-treated parasites (**Fig. 5D**). Thus, the vast majority of ATAC-seq peaks are diminished upon ACASi treatment (**Fig. 5E**).

Most of the 30 genes associated with ATAC-seq peaks only in ACASi-treated parasites are located in subtelomeric regions, and their transcripts are more abundant upon ACASi treatment (**Fig. 5E,F**, **Supp. Data 7**, **Supp. Data 9)**. Conversely, genes that were closest to ATAC-seq peaks that were shared between the two conditions or found only in the DMSO-treated parasites showed a significant decrease in transcript abundance upon ACASi treatment (**Fig. 5F**). In fact, there was a general decrease in ATAC-seq signal at the transcriptional start sites of all genes, regardless of whether their transcripts were more or less abundant upon ACASi treatment (**Fig. 5G**). These data demonstrate that while a small cohort of genes might have an increase in chromatin accessibility that leads to transcriptional upregulation upon ACASi treatment, most genes experience a loss of chromatin accessibility. These findings support a model in which chromatin accessibility and transcription is quickly shut down when acetyl-CoA is not readily available in the nucleus to facilitate histone acetylation.

## Discussion

In this study, we explored the nuclear role of acetyl-CoA synthetase in *P. falciparum*. Acetyl-CoA is essential to many cellular processes such as the TCA cycle, fatty acid synthesis, and acetylation of proteins such as histones. In *P. falciparum*, it is believed that while ACAS plays a role in acetyl-CoA production, the primary source of acetyl-CoA outside of the apicoplast is the mPDH (Cobbold *et al*, 2013; Nair *et al*, 2023). Interestingly, *Pf*ACAS and one subunit of the mPDH were shown to be essential at least during the asexual IDC, and both play a role in providing acetyl-CoA for histone acetylation (Zhang *et al*, 2018; Prata *et al*, 2021; Summers *et al*, 2022; Nair *et al*, 2023). However, we demonstrate that *Pf*ACAS also greatly influences chromatin composition and transcription.

While acetyl-CoA can diffuse through nuclear pores (Sivanand *et al*, 2018), a growing body of work has demonstrated specific roles for acetyl-CoA synthetases in the nucleus. It has been shown in multiple organisms that these synthetases associate with specific chromatin-associated complexes and genomic loci to provide local concentrated pools of acetyl-CoA to histone acetylases, which boost transcription of specific genes (Li *et al*, 2018). Nuclear translocation of acetyl-CoA synthetase enzymes in certain developmental or metabolic contexts leads to the activation of specific relevant cohorts of genes, linking cellular identity and milieu to a targeted transcriptional response. Such an enzyme would be an attractive target in the most virulent malaria parasite, *P. falciparum*.

We and others have shown that *Pf*ACAS can be present in both the cytoplasm and nucleus, and that inhibition or knockdown results in depletion of histone acetylation (Bryant *et al*, 2020; Prata *et al*, 2021; Summers *et al*, 2022; Nair *et al*, 2023). However, it was unclear 1) when and how *Pf*ACAS shuttles to the nucleus and 2) if *Pf*ACAS nuclear localization is required to generate acetyl-CoA in the nucleus for histone acetylation and transcriptional regulation. Regarding the first question, we found that while cytoplasmic levels of *Pf*ACAS remain fairly constant during the IDC, *Pf*ACAS becomes highly enriched in the nuclei of trophozoites, the most metabolically active stage of the IDC when the parasite undergoes rapid growth and acetyl-CoA levels are highest (Cobbold *et al*, 2016). It is possible that *Pf*ACAS is needed in the nucleus to generate acetyl-CoA especially at and after this time, which is also when DNA replication begins (Klaus *et al*, 2022; McDonald & Merrick, 2022; Castellano *et al*, 2024), total mRNA levels begin to increase substantially (Sims *et al*, 2009), and many histone residues become increasingly acetylated (Coetzee *et al*, 2017).

Our data suggest that nuclear translocation of *Pf*ACAS is responsive to cell cycle or metabolic cues. However, we were unable to separate the cytoplasmic and nuclear functions via preventing *Pf*ACAS entry into the nucleus. Our inability to mutate a conserved AMPK phosphorylation site in a putative C-terminal nuclear localization signal of *Pf*ACAS suggests its essentiality; however, treatment with salicylate, an AMPK activator used by (Mancio-Silva *et al*, 2017) to manipulate the divergent AMPK-like *Plasmodium* KIN kinase, did not affect *Pf*ACAS nuclear localization. It is possible that this site is phosphorylated by a different kinase or that *Pf*ACAS nuclear shuttling is controlled by a different pathway or post translational modifications. Future studies will help to discern the relative importance of the cytoplasmic versus nuclear roles of *Pf*ACAS.

Our findings provide new insight into how *Pf*ACAS inhibition affects histone acetylation and transcription. It has been suggested that mPDH is a major source of acetyl-CoA in the absence of acetate supplementation and provides acetyl-CoA for histone acetylation (Nair *et al*, 2023). However, our data suggest that *Pf*ACAS plays a more prominent role in histone acetylation than was previously thought. *Pf*ACAS inhibition for just three hours leads to a highly significant and global depletion of all acetylation sites on all histones assayed and withdrawal of *Pf*ACAS inhibitor for just 30 minutes restores histone acetylation levels to normal. This rapid response suggests that histone acetylation relies directly on nuclear *Pf*ACAS.

*Pf*ACAS inhibition also demonstrates the dynamic nature of histone acetylation in *P. falciparum*, suggesting a constant cycle of genome-wide acetylation and deacetylation to modulate genes during the rapid transcriptional cascade that drives the IDC. Indeed, recent genome-wide mapping of histone deacetylases SIR2A and HDAC1 revealed binding to many diverse cohorts of both transcriptionally repressed and active genes, potentially constantly producing a nuclear pool of acetate (Kanyal *et al*, 2024; Couble *et al*, 2025). As we and others (Prata *et al*, 2021) found *Pf*ACAS at low levels genome-wide, it could associate with chromatin to recycle acetate from histone deacetylases, providing acetyl-CoA to histone acetyltransferases in a constant cycle.

Conversely, high and specific enrichment at subtelomeric regions of *Pf*ACAS in association with a chromatin-remodeling/organizing complex could be particularly important to 1) balance the high acetate-generating histone deacetylation activity there in order to maintain a heterochromatic milieu and/or 2) boost acetylation of non-histone proteins such as AP2-P, which is highly enriched in the same region (Bryant *et al*, 2020) and was shown to be acetylated (Cobbold *et al*, 2016). Higher transcript abundance of *var* and *rifin* genes, which are located in subtelomeric regions, upon *Pf*ACAS inhibition might suggest that this enzyme plays a direct role in the transcriptional regulation of these genes; however, it is impossible to rule out if this change is simply due to disruption of the cell cycle.

Indeed, despite a global depletion of histone acetylation, *Pf*ACAS inhibition led to increases or decreases in transcript abundance for thousands of genes. This trend was also seen upon short-term treatment with an inhibitor of class I and II HDACs, trichostatin A, which led to elevated levels of histone acetylation (Andrews *et al*, 2012). These contradictory effects could be due to the deregulation of acetylation of non-histone proteins, such as transcription factors, chromatin remodelers, and even *Pf*ACAS itself (Miao *et al*, 2013; Cobbold *et al*, 2016), thereby amplifying the impact of *Pf*ACAS inhibition on chromatin architecture and gene expression. Regardless, we also observed a general reduction of chromatin accessibility after ACASi treatment, especially near transcriptional start sites, where histone acetylation has been shown to correlate with transcriptional activity of genes (Bártfai *et al*, 2010; Karmodiya *et al*, 2015). These data suggest a direct effect of loss of histone acetylation, leading to what we believe is a global transcriptional shut down and developmental arrest.

ACAS inhibition offers functional insight into the roles of histone acetylation beyond what is currently possible with inhibition or depletion of all histone acetyltransferases and provides evidence that histone acetylation is needed to maintain accessible, transcriptionally permissive chromatin. We propose that while mPDH may compensate for the loss of *Pf*ACAS activity in the cytoplasm, nuclear *Pf*ACAS is essential for histone acetylation dynamics in the fast-paced IDC. While it is still unclear whether *Pf*ACAS plays a role in directly regulating specific loci in the genome, its nuclear shuttling and important role in histone acetylation and global transcription warrant futures studies of how host metabolic status, and the fluctuation of host acetate levels especially, affects the transcriptional cascade that drives the parasite life cycle. Ultimately, this study adds to an ever-growing body of work that highlights the potential of transcriptional and epigenetic regulators as antiplasmodial targets.

## Materials and methods

### Parasite culture

Blood-stage 3D7 *P. falciparum* parasites were cultured in human RBCs supplemented with 2.5 g Albumax I (Thermo Fisher, 11020), hypoxanthine (0.1 mM final concentration, C.C.Pro Z-41-M) and 20 μg/ml gentamicin (Sigma, G1397) at 4% haematocrit and under 5% O_2_, 5% CO_2_ at 37°C. Parasites were synchronized by sorbitol lysis (5%, Sigma, S6021) at ring stage, followed by plasmagel (Plasmion, Fresenius Kabi) enrichment at late blood stages 24 h later. Another sorbitol treatment 6 h later, places the moment of re-invasion of the remaining parasites (or the 0 hours post invasion of the red blood cells) 3 h after the plasmagel enrichment. The resulting window of synchronicity is ±3 h for the cultures. Parasite development was monitored by Giemsa staining, and parasites were collected at 1-5% parasitemia. For **Figure 1E**, Gibco RPMI 1640 glucose-free (Thermo Fisher, 11879020) was used for Albumax I reconstitution and media preparation.

### Generation of strains

The ACAS-GFP-FKBP strain was generated using a two-plasmid system (pUF1 and pL7) based on the CRISPR/Cas9 system previously described (Ghorbal et al., 2014). A 3D7 wild-type bulk ring stage culture was transfected with 25 μg pUF1-Cas9 and 25 μg of pL7-PF3D7_ 0627800-GFP-ddFKBP containing a single guide RNA (sgRNA)-encoding sequence targeting the 3’ UTR of PF3D7_ 0627800 (**Supp. Data. 10**). The pL7-PF3D7_ 0627800-GFP-ddFKBP plasmid also contained a homology repair construct synthesized by GenScript Biotech (**Supp. Data 10**). This homology repair construct comprises a GFP-encoding and a destabilization domain FKBP-encoding sequence [L106P (Chu et al., 2008)], which are flanked by ∼300 bp homology repair regions upstream and downstream of the Cas9 cut site. One shield mutation was made in the upstream homology repair region to prevent further Cas9 cleavage of the modified locus. The sgRNA sequence was designed using Protospacer (MacPherson and Scherf, 2015). The sgRNA sequence uniquely targeted a single sequence in the genome. After transfection, drug selection was applied for five days at 2.67 nM WR99210 (Jacobus Pharmaceuticals) and 1.5 μM DSM1 (MR4/BEI Resources), and parasites were cultured in the constant presence of Shield-1 (AOB1848, Aobious) at 500 nM. Parasites reappeared approximately three weeks after transfection, and 5-fluorocytosine was used to negatively select the pL7 plasmid. Unfortunately, the FKBP destabilization domain did not respond to the presence or absence of Shield-1 (**Fig. EV1B**).

The ACAS-3HA strain was generated using the method of selection-linked integration (SLI) previously described (Birnbaum et al., 2017), with a slight modification wherein a GSG-encoding sequence (5’ – GGTAGTGGT – 3’) was added directly upstream of the T2A skip peptide-encoding sequence to enhance cleavage of the tagged protein from the downstream drug selection protein. The homology region corresponding to the 705 bp at the 3’ end of *acas* (PF3D7_0627800) was amplified using ACAS_3F/R (**Supp. Data 10**). The PCR fragment was then fused to a sequence encoding a 3HA epitope tag followed by a skip peptide followed by the neomycin resistance marker. A 3D7 wild-type bulk ring stage culture was transfected with 50 μg of pSLI plasmid (containing a yeast dihydroorotate dehydrogenase selection marker) and selected first with 1.5 μM DSM1 (MR4/BEI Resources). Parasites reappeared approximately three weeks after transfection and positive selection for integration was performed via the addition of 400µg/mL G418 (Sigma G8168).

The ACASΔCterm-3HA strain was generated by episomal transfection with pUF-ACASΔCterm-3HA. The PF3D7_0627800 coding sequence containing mutations in the 3’ end corresponding to the amino acid substitutions S946A, R951A, R952A (**Supp. Data 10**, GenScript Biotech) was fused to a 3HA-encoding sequence and inserted into the pUF plasmid under an *hsp90* promoter. A 3D7 wildtype clonal ring stage culture was transfected with 25 μg of pUF-ACASΔCterm-3HA (containing a yeast dihydroorotate dehydrogenase selection marker) and selected with 1.5 μM DSM1 (MR4/BEI Resources). Parasites reappeared approximately four weeks after transfection and were kept under DSM1 selection.

All cloning was performed using KAPA HiFi DNA Polymerase (Roche 07958846001), In-Fusion HD Cloning Kit (Clontech 639649), and XL10-Gold Ultracompetent E. coli (Agilent Technologies 200315). All the parasite lines were cloned by limiting dilution, and integration at the targeted genomic locus was confirmed by PCR (**Fig. EV1A, Supp. Data 10**) and Sanger sequencing.

### Cytoplasmic-nuclear fractionation and protein extraction

Synchronized parasite cultures at 2-3% parasitemia were centrifuged at 800 g for 3 min at 25°C. Medium was removed and the RBCs (1 ml) were lysed with 10 ml 0.075% saponin (Sigma S7900) in DPBS at 37°C. The parasites were centrifuged at 3,250 g for 3 min at 25°C and washed twice with 10 ml DPBS at 4°C. All further steps were performed at 4°C. The supernatant was removed, and the parasite pellet was resuspended in 900 μl of cytoplasmic lysis buffer [10 mM Tris–HCl pH 7.5, 10 mM NaCl, 1 mM EDTA, 0.65% IGEPAL CA-630 (Sigma), Complete EDTA-free protease inhibitors (PI, Roche 04693159001)] and rotated for 15 min. The lysates were centrifuged for 5 min at 2,000 *g* and the supernatant was collected as cytoplasmic fraction. The pellet was resuspended in 1 ml cytoplasmic wash buffer (10 mM Tris–HCl pH 7.5, 150 mM NaCl, 1 mM EDTA, PI) and rotated for 10 min. The lysates were centrifuged for 5 min at 5,000 *g*, and the supernatant was discarded. The pellet was resuspended in 200 μl of nuclear lysis buffer (10 mM Tris–HCl pH 7.5, 500 mM NaCl, 1 mM EDTA pH 8, 1% sodium deoxycholate, 1% IGEPAL, 0.1% SDS, PI), transferred to 1.5 ml sonication tubes (Diagenode C30010016), and sonicated for 2.5 min total (5 cycles of 30 s on/off) in a Diagenode Pico Bioruptor. The sonicated extracts were centrifuged at 13,500 *g* for 10 min, and the supernatant was collected as the nuclear fraction.

### Western blot analysis

Protein samples were supplemented with NuPage Sample Buffer (Thermo Fisher, NP0008) and NuPage Reducing Agent (Thermo Fisher, NP0004) and denatured for 10 min at 70 °C. Proteins were separated on a 4–12% Bis-Tris NuPage gel (Thermo Fisher, NP0321) and transferred to a nitrocellulose membrane with a Trans-Blot Turbo Transfer system (Bio-Rad). The membrane was blocked 1h at room temperature or overnight at 4°C with 1% milk in PBST (PBS, 0.1% Tween 20). HA-tagged proteins, GFP-tagged proteins, aldolase, histones, and histone PTMs were detected with anti-HA-HRP (Cell Signaling Technology C29F4, 1:1,000 in 1% milk-PBST), anti-GFP (ChromoTek PABG1,1:1,000 in 1% milk-PBST), anti-Aldolase-HRP (Abcam ab38905, 1:5,000 in 1% milk-PBST),) anti-H3 (Abcam ab1791, 1:2,500 in 1% milk-PBST), anti-H3K9ac (Millipore 07352, 1:1,000 in 1% milk-PBST), anti-H3 pan acetylation (Active Motif AM61638, 1:1,000 in 1% milk-PBST) and anti-H4 pan acetylation (Active Motif AM39026, 1:1,000 in 1% milk-PBST) primary antibodies, respectively. Anti-H3 or-GFP incubation was followed by donkey anti-rabbit secondary antibody conjugated to horseradish peroxidase (Cytiva NA934, 1:5,000 in 1% milk-PBST). HRP signal was developed with SuperSignal West Pico Plus chemiluminescent substrate (Thermo Fisher, 34580) for aldolase and H3 staining and SuperSignal West Femto maximum chemiluminescent substrate for HA and GFP staining (Thermo Fisher, 34096). Blots were imaged with a ChemiDoc XRS+ system (Bio-Rad) and saved using the Bio-Rad Image Lab software.

### Immunofluorescence Assay

Immunofluorescence assay was performed as in (Rosa *et al*, 2023). After blocking, fixed permeabilized cells were incubated overnight at 4 °C with anti-HA antibody (Roche 3F10, 1:500). After three PBS washes, cells were incubated with goat anti-rat Alexa Fluor 488 (Invitrogen, A-11006, 1:2,000) secondary antibody supplemented with DAPI (FluoProbes FP-CJF800, 20 μg/mL) for 30 min. Finally, cells were mounted with VectaShield (Vector Laboratories, H-1000) on glass slides. Images were acquired on a Deltavision Elite imaging system (GE Healthcare) and processed using the Icy software (de Chaumont *et al*, 2012).

### ACAS immunoprecipitation and mass spectrometry analysis

Sample preparation: Wildtype and ACAS-GFP parasites were grown and synchronized as described above. Five replicates of each strain containing 1.5×10^9^ parasites were harvested at 24 hpi and extracted from RBCs with saponin lysis, as described above. Each parasite pellet was resuspended in and crosslinked with 1 ml of 0.5 mM dithiobissuccinimidyl propionate in PBS (DSP, Thermo Fisher 22585) for 30 min at RT. The reaction was quenched by adding 6 ml of quenching buffer (25 mM Tris-HCl pH 7.5 in DPBS) and incubating for 10 min at RT. Samples were centrifuged at 1250 g at 4°C for 5 min, the supernatant was removed, and the pellet was washed with 10 ml DPBS at 4°C. The samples were centrifuged again, the supernatant was removed, and the pellets were snap frozen on dry ice and stored at-80°C. All samples were defrosted and processed as follows at 4°C. Cytoplasmic and nuclear fractions were made as described above, and each sample was incubated with 25 µl of Dynabeads Protein G (Invitrogen, 10004D) prebound to 10 µg anti-GFP antibody (Abcam, AB290) overnight at 4°C with gentle rotation. Dynabeads were isolated using a DynaMag magnet (Invitrogen) and washed three times with 200 µl wash buffer (10 mM Tris pH7.5, 0.5 mM EDTA, 150 mM NaCl and 0.05% IGEPAL), then resuspended in 100 µl of Ammonium Bicarbonate (25 mM). Proteins were reduced by adding 5 mM dithiothreitol (DTT) at pH 8.0 for 30 min at 57°C while shaking at 800 rpm. After cooling to room temperature, cysteines were alkylated by adding 10 mM iodoacetamide for 30 min in the dark. On bead-digestion was achieved overnight at 37°C by using 0.8 µg of Trypsin/Lys-C (Promega) while shaking at 800 rpm. Following digestion, samples were desalted using homemade C18 StageTips. Peptides were eluted using a ratio of 40/60 acetonitrile/water containing 0.1% formic acid and vacuum concentrated to dryness. Peptides were reconstituted in 10 µl of 0.3% trifluoroacetic acid (TFA) injection buffer before liquid chromatography-tandem mass spectrometry (LC-MS/MS) analysis.

LC-MS/MS: Online chromatography was performed with an RSLCnano LC system (UltiMate 3000, Thermo Scientific) coupled to an Orbitrap Eclipse mass spectrometer (Thermo Scientific). Peptides were trapped on a 2 cm nanoviper Precolumn (i.d. 75 μm, C18 Acclaim PepMap^TM^ 100, Thermo Scientific) at a flow rate of 3.0 µL/min in buffer A (2/98 acetonitrile/water containing 0.1% formic acid) for 4 min to desalt and concentrate the samples. Separation was performed on a 50 cm nanoviper column (i.d. 75 μm, C18, Acclaim PepMap^TM^ RSLC, 2 μm, 100Å, Thermo Scientific) regulated to a temperature of 50°C with a linear gradient from 2% to 25% buffer B (100% acetonitrile containing 0.1% formic acid) at a flow rate of 300 nL/min over 91 min. MS1 data were collected in the Orbitrap (120,000 resolution; maximum injection time 60 ms; Auto gain control (AGC) 4 x 10^5^). Charges states between 2 and 7 were required for MS2 analysis, and a 60 s dynamic exclusion window was used. MS2 scans were performed in the ion trap in rapid mode after fragmentation using higher-energy collisional dissociation (HCD: isolation window 1.2 Da; Normalized collision energy (NCE) 30%; maximum injection time 60 ms; AGC 10^4^).

Data processing: For identification, the data were searched against the *Plasmodium falciparum* 3D7 (UP000001450_36329) UniProt databases using Sequest HT through proteome discoverer (version 2.4). Enzyme specificity was set to trypsin and a maximum of two miss cleavages sites were allowed. CAMthiopropanoyl lysine, Oxidized methionine, Met-loss, Met-loss-Acetyl and N-terminal acetylation were set as variable modifications. Carbamidomethylation of cysteins were set as fixed modification. Maximum allowed mass deviation was set to 10 ppm for monoisotopic precursor ions and 0.6 Da for MS/MS peaks. The resulting files were further processed using myProMS v3.10.0 (https://github.com/bioinfo-pf-curie/myproms) (Poullet *et al*, 2007). FDR calculation used Percolator (The *et al*, 2016) and was set to 1% at the peptide level for the whole study. The label free quantification was performed by peptide Extracted Ion Chromatograms (XICs), reextracted across all conditions and computed with MassChroQ version 2.2.21(Valot *et al*, 2011). For protein quantification, XICs from proteotypic peptides shared between compared conditions (TopN) and missed cleavages were allowed. Median and scale normalization was applied on the total signal to correct the XICs for each biological replicate (N=5 in each condition). To estimate the significance of the change in protein abundance, a linear model (adjusted on peptides and biological replicates) was performed, and *p*-values were adjusted using the Benjamini–Hochberg FDR procedure. Proteins with at least 3 total peptides in all states and an adjusted p-value ≤ 0.05 were considered significantly enriched in sample comparisons. Proteins unique to a condition were also considered if they matched the peptides criteria.

### NLS conservation analysis

Putative C-terminal AMPK phosphorylation sites of *Hs*ACSS2 and *Pf*ACAS were aligned using Clustal Omega (Madeira *et al*, 2024).

### Immunoprecipitation and western blot of ACAS

Parasites were grown and synchronized as described above. 2 ml of iRBCs at 2-3% parasitemia were collected at 24 hpi and lysed with saponin as described above. Cytoplasmic and nuclear fractions were made as described above with PI and phosphatase inhibitor (PhosphoSTOP, Roche, 4906845001). Immunoprecipitation was performed as described previously (Rosa *et al*, 2023) Briefly; the entire cytoplasmic or nuclear fraction was incubated overnight at 4°C with gentle rotation with 25 µl Protein G Dynabeads (10004D, Invitrogen) bound to 2 µg anti-HA antibody (Abcam, ab9110). Beads were isolated using a DynaMag magnet, and supernatant was removed. Beads were washed three times with 200 µl wash buffer (10mM Tris pH7.5, 0.5mM EDTA, 150mM NaCl, 0.05% IGEPAL) at 4°C, then resuspended in sample buffer with reducing agent and incubated for 10 mins at 70°C. Western blot analysis was performed as described above using one third of the final IP sample for the anti-HA (1:1000 in 1% milk-PBST, Abcam ab9110) blot, and two thirds for the blot that was probed with anti-Phospho Ser/Thr (1:1000 in 1% milk-PBST Abcam ab17464), anti-aldolase-HRP (AB38905), and anti-H3 (Abcam ab1791) primary antibodies.

### Chromatin immunoprecipitation and sequencing

Clonal populations of ACAS-3HA and ACAS-GFP-FKBP parasites (1.36 10^9^ and 8 x 10^8^ parasites respectively) were tightly synchronized and collected at 24 hpi. In each case, parasite culture was centrifuged at 800 g for 3 min at 25°C. Medium was removed, and the RBCs were lysed with 10 ml 0.075% saponin (Sigma, S7900) in DPBS at 37°C. The parasites were centrifuged at 3,250 g for 3 min at 25°C and washed with 10 ml DPBS at 37°C. For the ACAS-GFP-FKBP parasites, the supernatant was removed, and the parasite pellet was resuspended in 10 ml PBS at 25°C. The parasites were cross-linked by adding methanol-free formaldehyde (Thermo Fisher, 28908) (final concentration 1%) and incubating with gentle agitation for 10 min at 25°C. The cross-linking reaction was quenched by adding glycine (final concentration 125 mM; Sigma, G8899) and incubating with gentle agitation for 5 min at 25°C. Parasites were centrifuged at 3,250 g for 5 min at 4°C and the supernatant removed. Parasite pellets were snap frozen and stored at −80°C. For the ACAS-3HA parasites after saponin lysis, the supernatant was removed, and the parasite pellet was resuspended in 20 ml of PBS at 25°C. MgCl_2_ (Invitrogen, AM9530G) was added to a final concentration of 1 mM. A volume of 160 μl of ChIP Cross-link Gold (Diagenode, C01019027) was added, and the sample was incubated at 25°C for 30 min with gentle agitation. Parasites were centrifuged at 3,250 g for 5 min at 4°C and the supernatant removed. The pellet was washed twice with DPBS at 4°C and centrifuged at 3,250 g for 5 min at 4°C. The parasites were resuspended in 20 ml DPBS at 4 °C and were further crosslinked by adding methanol-free formaldehyde (Thermo Fisher, 28908) (final concentration 1%) and incubating with gentle agitation for 15 min at 25°C. The cross-linking reaction was quenched by adding glycine (final concentration 125 mM; Sigma, G8899) and incubating with gentle agitation for 5 min at 25 °C. Parasites were centrifuged at 3,250 g for 5 min at 4°C, and the supernatant removed. Parasite pellets were snap frozen and stored at −80 °C.

For each ChIP, 200 µl of Protein G Dynabeads (Invitrogen, 10004D) were washed twice with 1 ml ChIP dilution buffer (16.7 mM Tris-HCl pH 8, 150 mM NaCl, 1.2 mM EDTA pH 8, 1% Triton X-100, 0.01% SDS) using a DynaMag magnet (Thermo Fisher, 12321D). The beads were resuspended in 1 ml ChIP dilution buffer with 8 μg of anti-HA antibody (Abcam, ab9110) or 40 μg anti-GFP antibody (Abcam ab290) and incubated on a rotator at 4°C for 6 h.

The cross-linked parasites were resuspended in 4 ml of lysis buffer (10 mM HEPES pH 8, 10 mM KCl, 0.1 mM EDTA pH 8, PI) at 4°C, and 10% IGEPAL CA-630 (Sigma) was added (final concentration 0.25%). The parasites were lysed in a prechilled dounce homogenizer (100 strokes). The lysates were centrifuged for 10 min at 13,500 g at 4 °C, the supernatant was removed, and the pellet was resuspended in 3.6 ml SDS lysis buffer (50 mM Tris-HCl pH 8, 10 mM EDTA pH 8, 1% SDS, PI) at 4°C. The liquid was distributed into 1.5 ml sonication tubes (Diagenode, C30010016, 300 µl per tube) and sonicated for 12 min total (24 cycles of 30 s on/off) in a Diagenode Pico Bioruptor at 4°C. The sonicated extracts were centrifuged at 13,500 g for 10 min at 4°C, and the supernatant, corresponding to the chromatin fraction, was kept. The DNA concentration was determined using the Qubit dsDNA High Sensitivity Assay kit (Thermo Fisher, Q32851) with a Qubit 3.0 fluorometer (Thermo Fisher). Chromatin lysate corresponding to 100 ng of DNA was diluted in SDS lysis buffer (final volume 200 μl) and kept as ‘input’ at −20°C. Chromatin lysate corresponding to 8 μg (for ACAS-3HA) and 4.588 μg (for ACAS-GFP-FKBP) of DNA was diluted 1:10 in ChIP dilution buffer at 4°C.

Using a DynaMag magnet, the antibody-conjugated Dynabeads were washed twice with 1 ml ChIP dilution buffer and resuspended in 100 μl of ChIP dilution buffer at 4°C. Then the washed antibody-conjugated Dynabeads were added to the diluted chromatin sample and incubated overnight with rotation at 4°C. The beads were collected on a DynaMag into 8 different tubes per sample, the supernatant was removed, and the beads in each tube were washed for 5 min with gentle rotation with 1 ml of the following buffers, sequentially:

○Low-salt wash buffer (20 mM Tris-HCl pH 8, 150 mM NaCl, 2 mM EDTA pH 8, 1% Triton X-100, 0.1% SDS) at 4°C.

○High-salt wash buffer (20 mM Tris-HCl pH 8, 500 mM NaCl, 2 mM EDTA pH 8, 1% Triton X-100, 0.1% SDS) at 4°C.

○LiCl wash buffer (10 mM Tris-HCl pH 8, 250 mM LiCl, 1 mM EDTA pH 8, 0.5% IGEPAL CA-630, 0.5% sodium deoxycholate) at 4°C.

○TE wash buffer (10 mM Tris-HCl pH 8, 1 mM EDTA pH 8) at 25 °C.

After the washes, the beads were collected on a DynaMag, the supernatant was removed, and the beads were resuspended in 800 μl (100 μl/tube) of elution buffer and incubated at 65°C for 30 min with agitation (1,000 rpm 30 s on/off). The beads were collected on a DynaMag and the eluate, corresponding to the ‘ChIP’ samples, was transferred to a different tube (200 μl/tube).

For purification of the DNA, both ‘ChIP’ and ‘Input’ samples were incubated for ∼10 h at 65°C to reverse the crosslinking. 200 μl TE buffer followed by 8 μl of RNase A (Thermo Fisher, EN0531) (final concentration of 0.2 mg/ml) were added to each tube, which was then incubated for 2 h at 37°C. 4 μl Proteinase K (New England Biolabs, P8107S) (final concentration of 0.2 mg/ml) was added to each tube, which was then incubated for 2 h at 55°C. A volume of 400 μl phenol:chloroform:isoamyl alcohol (25:24:1) (Sigma, 77617) was added to each tube, which was then mixed by vortexing and centrifuged for 10 min at 13,500 g at 4°C to separate phases. The aqueous top layer was transferred to another tube and mixed with 30 μg glycogen (Thermo Fisher, 10814) and 5 M NaCl (200 M final concentration). Ethanol (800 μl, 100%) at 4°C was added to each sample, which was then incubated at −20°C for 30 min. The DNA was pelleted by centrifugation for 10 min at 13,500 g at 4°C, washed with 500 μl 80% ethanol at 4°C, and centrifuged for 5 min at 13,500 g at 4°C. After removing the ethanol, the pellet was dried at 25°C and all DNA for each sample (ChIP and Input) was resuspended in 30 μl 10 mM Tris-HCl pH 8. The DNA concentration and average size of the sonicated fragments were determined using a DNA high sensitivity kit and an Agilent 2100 Bioanalyzer. Libraries for Illumina next generation sequencing were prepared using the MicroPlex library preparation kit (Diagenode, C05010014), with KAPA HiFi polymerase (KAPA Biosystems) substituted for the PCR amplification. Libraries were sequenced on the NextSeq 500 platform (Illumina).

### ChIP-seq processing and analysis

Sequenced reads (150 bp paired end) were mapped to the *P. falciparum* genome (plasmoDB.org, v.3, release 56) using Bowtie2 (Langmead & Salzberg, 2012). PCR duplicates were filtered using samtools’ fixmate and markdup commands (Li *et al*, 2009), and only alignments with a mapping quality ≥30 were retained (samtools view-q 30) (Li *et al*, 2009). The paired-end deduplicated ChIP and input BAM files were used as treatment and control, respectively, for peak calling with the MACS3 “macs3 callpeak-g 23e6-q 0.001” command (Zhang *et al*, 2008). The “--nomodel” flag was added when necessary. For ACAS-3HA and ACAS-GFP-FKBP samples, consensus peaks were defined using the bedtools intersect command (Quinlan & Hall, 2010). ChIP/input ratio tracks were generated using deeptool’s bamCompare command (Ramírez *et al*, 2016). Integrative Genomics Viewer was used to inspect tracks and MACS3 peaks (Robinson *et al*, 2011). Binding peaks were associated with the nearest protein-coding genes using bedtools closest command along with *P. falciparum* reference genome feature file (gff) (plasmoDB.org, v.3, release 56). Only regions 500 bp upstream or downstream of the protein-coding genes were considered further for downstream analysis. To perform Gene Ontology Enrichment analysis on the genes closest to the peaks, GO Enrichment tool from PlasmoDB web interface (plasmoDB.org, v.3, release 56) was used for Ontology Term “Biological Process”, with a P value cut-off of 0.05. Plotting was performed in R using the ChIPseeker (Yu *et al*, 2015) and karyoplotter (Gel & Serra, 2017) packages.

### Histone purification and mass spectrometry

3D7 wildtype parasites were grown and synchronized as described above. Six replicates of 2 ml of iRBCs each at 2.5% parasitemia at 21 hpi were incubated with 2.5 µM ACAS inhibitor [MMV019721, EC_50_ = 460nM (Summers *et al*, 2022)] or DMSO (Sigma, D8418) for 3 h. At 24 hpi, parasites were extracted from RBCs with saponin lysis, as described above, and the pellet was snap frozen and stored at-80°C. Histones were extracted as previously described (Couble *et al*, 2025). The final histone pellets were allowed to dry for 10 min under a chemical hood and then were immediately stored at-80°C. Samples were digested using trypsin in a 1:50 ratio (protease:protein) in 100 mM ammoniumbicarbonate supplemented with 5 mM Tris(2-carboxyethyl)phosphine hydrochloride (TCEP) and 20 mM 2-chloroacetamide (CAA). Digestion was carried out overnight at 37°C and stopped the next day by the addition of TFA to 1%. Samples were desalted using Oasis HLB µElution Plates (Waters), according to the manufacturer’s recommended protocol.

An UltiMate 3000 RSLCnano LC system equipped with a trapping cartridge (µ-Precolumn C18 PepMap™ 100, 300 µm i.d. × 5 mm, 5 µm particle size, 100 Å pore size; Thermo Fisher Scientific) and an analytical column (nanoEase™ M/Z HSS T3, 75 µm i.d. × 250 mm, 1.8 µm particle size, 100 Å pore size; Waters) was used. Samples were trapped at a constant flow rate of 30 µl/min using 0.05% trifluoroacetic acid (TFA) in water for 6 min. After switching in-line with the analytical column, which was pre-equilibrated with solvent A (3% DMSO, 0.1% formic acid in water), the peptides were eluted at a constant flow rate of 0.3 µl/min using a gradient of increasing solvent B concentration (3% DMSO, 0.1% formic acid in acetonitrile). Peptides were introduced into the Orbitrap Fusion™ Lumos™ Tribrid™ mass spectrometer (Thermo Fisher Scientific) via a Pico-Tip emitter (360 µm OD × 20 µm ID; 10 µm tip, CoAnn Technologies) using an applied spray voltage of 2.4 kV. The instrument was operated in positive ion mode, and the capillary temperature was set to 275°C. Full MS scans were acquired in profile mode over an m/z range of 350–2000, with a resolution of 120,000 at m/z 200 in the Orbitrap. The maximum injection time was set to 50 ms, and the AGC target was set to ‘standard’. The instrument was operated in data-dependent acquisition (DDA) mode, with MS/MS scans acquired in the Orbitrap at a resolution of 15,000. Fixed first mass was set to 110 m/z. The maximum injection time was set to 54 ms, and the AGC target was set to 400%. Fragmentation was performed using HCD with a normalized collision energy of 34%. MS2 data were acquired in profile mode, and dynamic exclusion was set to 30 sec.

Raw files were then searched using MaxQuant (version 2.4.9.0) (Cox & Mann, 2008) against the FASTA databases UP000001450 (*Plasmodium falciparum*, isolate 3D7, ID36329, 5372 entries, March 2023), and the sequences of the *Plasmodium falciparum* H2A, H2AZ, H2B, H2BZ, H3, H3.3 and H4 histones. The following modifications were included into the search parameters: Carbamidomethylation on C as fixed modification; Oxidation (M) and Acetylation (protein N-terminus and K) as variable modifications. In the protein quantification, only unmodified and oxidated as well as protein N-terminus peptides were included. For the full scan (MS1) and the MS/MS (MS2) spectra a mass error tolerance of 20 PPM was set. For protein digestion,’trypsin’ was used as protease with an allowance of maximum 2 missed cleavages requiring a minimum peptide length of 7 amino acids. Match between runs was enabled, as well as IBAQ value calculation. The false discovery rate on peptide and protein level was set to 0.01. MS1 filtering was done in Skyline 23.1.0 (Schilling *et al*, 2012). The area under the curve values of the extracted ion chromatograms of the identified acetylated peptides were exported and normalized with the input of the respective protein. The data was log2 transformed and fold changes and *p*-values were calculated by same variance bi-directional Student’s t-test.

### Mass spectrometry-based lipidomic analyses

Wildtype parasites were synchronized as described above, and the culture was split into eight flasks at 21 hpi. Four flasks were treated with 2.5 µM MMV019721 *Pf*ACAS inhibitor, and the other four received an identical volume of DMSO (Sigma, D8418). Cultures were incubated for 3 h, then RBCs were lysed in 0.075% saponin (Sigma, S7900) in PBS at 25°C. Samples were centrifuged at 3,250 g for 5 min at 4°C, washed in PBS at 4°C, centrifuged at 3,250 g for 5 min at 4°C, and the pellets were directly stored at-80 °C.

Analyses were performed as previously described and modified for *P. falciparum* (Huynh *et al*, 2019; Arnold *et al*, 2025; Dass *et al*, 2024). Briefly, parasites were metabolically quenched in dry ice - ethanol for one min then washed three times in cold phosphate-buffered saline. Total lipid was extracted using 350 µL chloroform/methanol (4:3 v/v) containing 1 nmol each of internal standards (Avanti-Sigma), followed by the addition of 140 µL of water to induce biphasic separation. The lower organic phase was collected and subsequently dried in a vial insert. For Total FA analysis by GCMS, pellets were washed 3x times in cold PBS to eliminate lipids from the culture media before conservation at-80°C. Internal standards 10 µmol FFA C13:0 and 10 µmol PC C21:0 (Avanti Polar lipids) were added to samples, and lipids were extracted following the Bligh and Dyer modified by Folch protocol (Ramakrishnan *et al*, 2012). FAs were derivatized using trimethylsulfonium hydroxide (TMSH) and analyzed by gas-chromatography mass-spectrometry (Agilent 5977A-7890B) (GC-MS). Fatty acid methyl esters were identified by their mass spectrum and retention time compared to authentic standards. For total lipid analysis, the recovered lipids were dissolved in methanol (50 µL) and an aliquot (10 µL) was injected into an Agilent 1290 infinity/Infinity II LCMS system equipped with ZORBAX Eclipse Plus C18, 100 x 2.1 mm, 1.8 µm reversed-phase column (Agilent) with Infinity II inline filter, 0.3 um (Agilent) maintained at 45°C. Samples were separated by the change in gradient of solvent A [qater:acetonitrile:isopropanol, 5:3:2 (v/v/v), 10 mM ammonium formate] and solvent B [isopropanol:acetonitrile:water, 90:9:1 (v/v/v), 10 mM ammonium formate]. The gradient was as follows, starting with a flow rate of 0.4 mL/min at 15% B and increasing to 50% over 2.5 min, then to 57% at 2.6 min, then to 70% over 9 min, and to 93% at 9.1 min, then to 96% over 11 min, and to 100% at 11.1 min and hold to 12 min then back to 15% at 12.2 min to 16 min (total of 16 min). MS (Agilent 6495c triple quadrupole) was operated in for targeted analysis with DMRM (dynamic multiple reaction monitoring) with 650 ms of cycle time with the following setting: Agilent Jet stream ion source with positive and negative switching with gas temperature at 150°C, drying gas (nitrogen) 17 L/min, nebulizer gas 20 psi, sheath gas temperature at 200°C with flow at 10 L/min, capillary voltage 3500 V for positive and-3000 V for negative mode, nozzle voltage at 1000 V for positive and-1500 V for negative mode. LCMS data was combined to pre-existing dynamic multiple reaction monitoring (DMRM) method for optimized targeted lipidomic analysis for *P. falciparum*.

### RNA extraction and mRNA sequencing

Wildtype parasites were synchronized as described above, and the culture was split into six flasks at 21 hpi. Three flasks were treated with 2.5 µM MMV019721 *Pf*ACAS inhibitor, and the other three received an identical volume of DMSO (Sigma, D8418). Cultures were incubated for 3 h, then RBCs were lysed in 0.075% saponin (Sigma, S7900) in PBS at 37°C. Samples were centrifuged at 3,250 g for 5 min, washed in PBS at 37°C, centrifuged at 3,250 g for 5 min, and resuspended in 700 μl QIAzol reagent (Qiagen, 79306). RNA was extracted using an miRNeasy Mini kit (Qiagen, 1038703) with the recommended on-column DNase treatment. Total RNA was poly(A) selected using the Dynabeads mRNA Purification kit (Thermo Fisher, 61006). Library preparation was performed using the NEBNext Ultra II Directional RNA Library Prep Kit for Illumina (New England Biolabs, E7760S) and paired-end sequencing was performed on the Nextseq 500 platform (Illumina).

### RNA-seq processing and analysis

Sequenced reads were mapped to the *P. falciparum* genome (plasmoDB.org, v.3, release 56) using ‘STAR’ (Dobin *et al*, 2013), restricting the number of multiple alignments allowed for a read using the option ‘–outFilterMultimapNmax 1’. Alignments were subsequently filtered for duplicates and a mapping quality ≥20 using samtools (Li *et al*, 2009). Gene counts were quantified with htseq-count (Anders *et al*, 2015) and differentially expressed genes were identified in R using the package DESeq2 (Love *et al*, 2014). Significantly differentially expressed genes were identified as those with a Benjamini–Hochberg-adjusted P-value (i.e., q) ≤ 0.05. MA plots were generated using the “baseMean” (mean normalized read count over all replicates and conditions) and “log2FoldChange” values (ACAS inhibitor-treated over DMSO control) as determined by DESeq2. RPKM values were calculated in R using the command rpkm() from the package edgeR (Robinson *et al*, 2010). Gene Ontology enrichment analysis was performed on significantly differentially expressed genes (q < 0.05) with over two-fold change using the built-in tool at PlasmoDB.org (Aurrecoechea *et al*, 2017) (v.3, release 56) with default settings for Biological Process (P < 0.05).

### Estimation of cell cycle progression

RNA-seq-based cell cycle progression was estimated in R by comparing the normalized expression values (i.e., RPKM, reads per kilobase per exon per one million mapped reads) of each sample to the microarray data from (Bozdech *et al*, 2003) using the statistical model as in (Lemieux *et al*, 2009).

### ATAC sequencing

Wildtype parasites were synchronized as described above, and the culture was split into four flasks at 21 hpi. Two flasks were treated with 2.5 µM MMV019721 *Pf*ACAS inhibitor, and the other two received an identical volume of DMSO (Sigma, D8418). Cultures were incubated for 3 h, then RBCs (equivalent to 5×10^6^ parasites for each replicate) were lysed in 0.075% saponin (Sigma, S7900) in PBS at 37°C. Samples were centrifuged at 3,250 g for 5 min, washed in PBS at 37°C, centrifuged at 3,250 g for 5 min, and resuspended in 500 µL of cold PBS containing protease inhibitors from the Diagenode ATAC-seq kit (Hologic Diagenode Cat# C01080002). Tubes were centrifuged at 3250 g for 5 min, and supernatant was removed. ATAC-seq was performed following the manufacturer’s instructions. Nuclei were extracted by adding 50 µL of cold ATAC Lysis Buffer 1 containing 2% Digitonin to the cell pellet for 3 min. Nuclei were collected by centrifugation, and a transposition reaction was performed by adding 50 µL of tagmentation solution (containing Tagmentation buffer, Tagmentase, 2% Digitonin, 10% Tween20, PBS and Nuclease-free water) for 30 min at 37°C. DNA was isolated using supplied spin columns and eluted in 12 µL of DNA elution buffer. Tagmented DNA was amplified using UDI for Tagmented libraries (Diagenode; #C01011035), and libraries were purified using AMPure XP beads (Beckman Coulter; #A63881). Quality and size distribution assessment of the libraries was performed on a Bioanalyzer 2100 system (Agilent) and by qPCR on a CFX384 real-time PCR detection system (Bio-Rad) using the kit’s qPCR mix and QC guidelines. Sequencing was performed on the Nextseq 2000 platform (Illumina).

### ATAC-seq processing and analysis

Sequenced reads (150 bp paired end) were analyzed using the nf-core ATAC-seq pipeline dev (https://nf-co.re/atacseq/2.1.2/) The pipeline was run on all ATAC-seq samples, including the 25h timepoint dataset from (Toenhake *et al*, 2018), using the *P. falciparum* genome and annotation files (plasmoDB.org, v.3, release 56), masked for rDNA loci. Notable parameters used were: For the Bowtie2 (Langmead & Salzberg, 2012) step, external arguments “-I 50-X 150” to only keep fragments with a size between 50 and 150 bp following guidelines and methodology from (Toenhake *et al*, 2018).

For the MACS3 (Zhang *et al*, 2008) peak calling step, *P. falciparum* gDNA ATAC-seq data from (Toenhake *et al*, 2018) was used as control with the *--keep_mito --macs_gsize 23000000--macs_fdr 0. 001*. Finally, for consensus peak identification, parameter “*--min_reps_consensus 2”* was specified to the pipeline to only keep peaks present in at least the two replicates in a given condition. Plotting of Violin plots was performed in R using the ggplot2 package (Wickham, 2016).

## Supplementary Data Legends

**Supplementary Data 1:** Label-free quantitative LC-MS/MS analysis of proteins immunoprecipitated with GFP antibody in *Pf*ACAS-GFP versus wildtype trophozoite cytoplasmic extracts. *Pf*ACAS (PF3D7_0627800) is highlighted in grey. Tab 1: all *P. falciparum* proteins detected. Tab 2: all *P. falciparum* proteins detected with adjusted *p*-value < 0.05. Tab 3: all *P. falciparum* proteins detected with adjusted *p*-value < 0.05 and ≥ 3 peptides in all replicates.

**Supplementary Data 2:** Label-free quantitative LC-MS/MS analysis of proteins immunoprecipitated with GFP antibody in *Pf*ACAS-GFP versus wildtype trophozoite nuclear extracts. *Pf*ACAS (PF3D7_0627800) is highlighted in grey. Tab 1: all *P. falciparum* proteins detected. Tab 2: all *P. falciparum* proteins detected with adjusted *p*-value < 0.05. Tab 3: all *P. falciparum* proteins detected with adjusted *p*-value < 0.05 and ≥ 3 peptides in all replicates.

**Supplementary Data 3**: Gene Ontology analysis (biological process) for proteins that are significantly enriched in the cytoplasmic (tab 1) or nuclear (tab 2) IP/MS of *Pf*ACAS-GFP trophozoites (defined in **Supp. Data 1,2**). Shown are Gene Ontology ID, name of category, total number of genes in this category (Bgd count), number of genes from the query in this category (Result count), genes from the query in this category (Result gene list), percentage of genes from the query in this category out of total number of genes in this category (Pct of bgd), P value, and Bonferroni adjusted P value.

**Supplementary Data 4**: Peaks from ACAS-3HA and ACAS-GFP ChIP-seq experiments in trophozoites and their chromosomal coordinates, summit location relative to peak start, score [-log10(q-value)], fold enrichment (FE) at the summit, P value, q-value, the closest unique protein-coding gene and its product description. Tab 1: consensus peaks. Tab 2: ACAS-GFP peaks. Tab 3: ACAS-3HA peaks.

**Supplementary Data 5:** Mass spectrometry data for histones extracted from wildtype trophozoites (24 hpi) treated for three hours with DMSO or ACASi.

Tab 1: Acetylated peptides identified with Maxquant on histone proteins

Tab 2: Total MS1 area values (abundance of acetylated peptides) after MS1 filtering exported from the Skyline software

Tab 3: Total protein abundances for histone proteins quantified by MaxQuant

Tab 4: Original MaxQuant output file “acetyl (K)Sites.txt” Tab 5: Original MaxQuant output file “proteinGroups.txt”

**Supplementary Data 6:** Lipidomics GC/MS TMSH quantification. Shown are nM of lipids (tab 1) or fatty acids (tab 2) per 10^7^ parasites for replicates of wildtype trophozoites (24 hpi) treated for three hours with DMSO or ACASi. Lipid abbreviations: CE: cholesteryl-ester, COH: cholesterol, MAG: monoacylglycerol, DAG: diacylglycerol, TAG: triacylglycerol, Cer: ceramide, FFA: free fatty acid, LPC: lyso-phosphatidylcholine, LPE: lyso-phosphatidylethanolamine, PC: phosphatidylcholine, PE: phosphatidylethanolamine, PE-P: phosphatidylethanolamine plasmalogen, PE-Cer: phosphatidylethanolamine ceramide, PG: phosphoglycerol, PI: phosphatidylinositol, PS: phosphatidylserine, SM: sphingomyelin, PA: phosphatidic acid, CL: cardiolipin. Fatty acid nomenclature of CXX:Y (XX: number of carbons, Y: number of insaturations).

**Supplementary Data 7**: Differential gene expression analysis in trophozoites of ACAS inhibitor-treated over DMSO-treated wildtype parasites. Tab 1: all genes. Tab 2: genes whose transcripts are significantly at least two-fold more abundant upon ACASi treatment. Tab 3: genes whose transcripts are significantly at least two-fold less abundant upon ACASi treatment. Tab 4: *var* and *rifin* genes.

**Supplementary Data 8:** Gene Ontology analysis (biological process) for genes whose transcripts are significantly at least two-fold more (tab 1) or less (tab 2) abundant upon ACAS inhibitor treatment in trophozoites (defined in **Supp. Data 7**). Shown are Gene Ontology ID, name of category, total number of genes in this category (Bgd count), number of genes from the query in this category (Result count), genes from the query in this category (Result gene list), percentage of genes from the query in this category out of total number of genes in this category (Pct of bgd), P value, and Bonferroni adjusted P value.

**Supplementary Data 9:** Consensus peak intervals from ACASi-and DMSO-treated replicates from ATAC-seq experiment in trophozoites (24 hpi). Shown for each interval are the chromosome coordinates, ID, nearest feature, nearest gene, and distance to the nearest transcriptional start site. Shown for each interval are the number of peaks included, the number of samples in which it is present, its presence in each replicate, and its fold enrichment (relative to gDNA control), log_10_(p-value), and log_10_(q-value) for each replicate.

**Supplementary Data 10:** Primers, oligos, and DNA blocks used in this study

## Data availability

ChIP-seq, RNA-seq, and ATAC-seq datasets generated in this study are available on NCBI with BioProject accession PRJNA1477376. ACAS IP-MS and histone MS raw data generated in this study are available on the ProteomeXchange Consortium via the PRIDE (Perez-Riverol *et al*, 2025) partner repository with the dataset identifiers PXD079912 (reviewer password: A80gLWOPppD6) and PXD080332 (reviewer password: DMBS0tzL5SE7), respectively.

## Author contributions

J.M.B. conceptualized the project. R.F.A., P.C., J.J.S., F.D., C.R., G.L.M., Y.Y.B., and J.M.B. performed experiments. R.F.A., J.J.S., F.D., Y.Y.B., J.M.B., and S.B. conducted formal analysis. R.F.A., P.C., J.J.S., Y.Y.B., and J.M.B. prepared data visualizations. J.M.B., R.F.A., and P.C. wrote the original draft of the paper. All authors reviewed and edited the paper. J.M.B., D.L., and C.B. acquired funding. J.M.B., S.B., D.L., C.B., and A.S. supervised the project.

## Conflict of interest

The authors declare that they have no conflict of interest.

## Acknowledgements

J.M.B. and her group were supported by the French Agence Nationale de la Recherche (ANR-21-CE15-0002-02, ANR-21-CE15-0010-01.) Work at the Institut Curie LSMP was supported by “la Région Île-de-France” (EX061034) and ITMO Cancer of Aviesan and INCa on funds administered by Inserm (21CQ016-00). C.Y.B., Y.Y.B., and the GEMELI lipidomics platform were supported by the French Agence Nationale de la Recherche (ANR-21-CE44-0010, ANR-23-CE15-0009-01, ANR-24-CE15-2171-02, ANR-25-CE30), Fondation pour la Recherche Médicale (FRM EQU202103012700), Laboratoire d’Excellence (LabEx) Parafrap, France (ANR-11-LABX-0024), LIA-IRP CNRS Program (Apicolipid project), the Université Grenoble Alpes (IDEX ISP Apicolipid), Région Auvergne Rhone-Alpes (Grant IRICE Project GEMELI), the Collaborative Research Program Grants CEFIPRA (MESRI-DBT, Project 6003-1, IARDP grant #2023-0414), and IBiSA for the Gemeli Platform. S.B. was supported by an ERC Starting Grant (947819). The LSMP thanks Patrick Poullet (bioinformatics platform Institut Curie U1331) for the continuous development of the myProMS software. The members of the Plasmodium Chromatin and Transcription Group thank Dr. Artur Scherf for his invaluable support and guidance, Dr. Beatriz Baragana (University of Dundee) for kindly providing the drug MMV019721, and Drs. Manuel Llinás and Isadora Prata for their critical reading of and helpful feedback on this manuscript. We acknowledge the use of the Biomics, PBI, and PFC platforms at the Institut Pasteur, the support of the LabEx ParaFrap network, and the essential resource PlasmoDB.

---

**Expanded View Figure 1:**
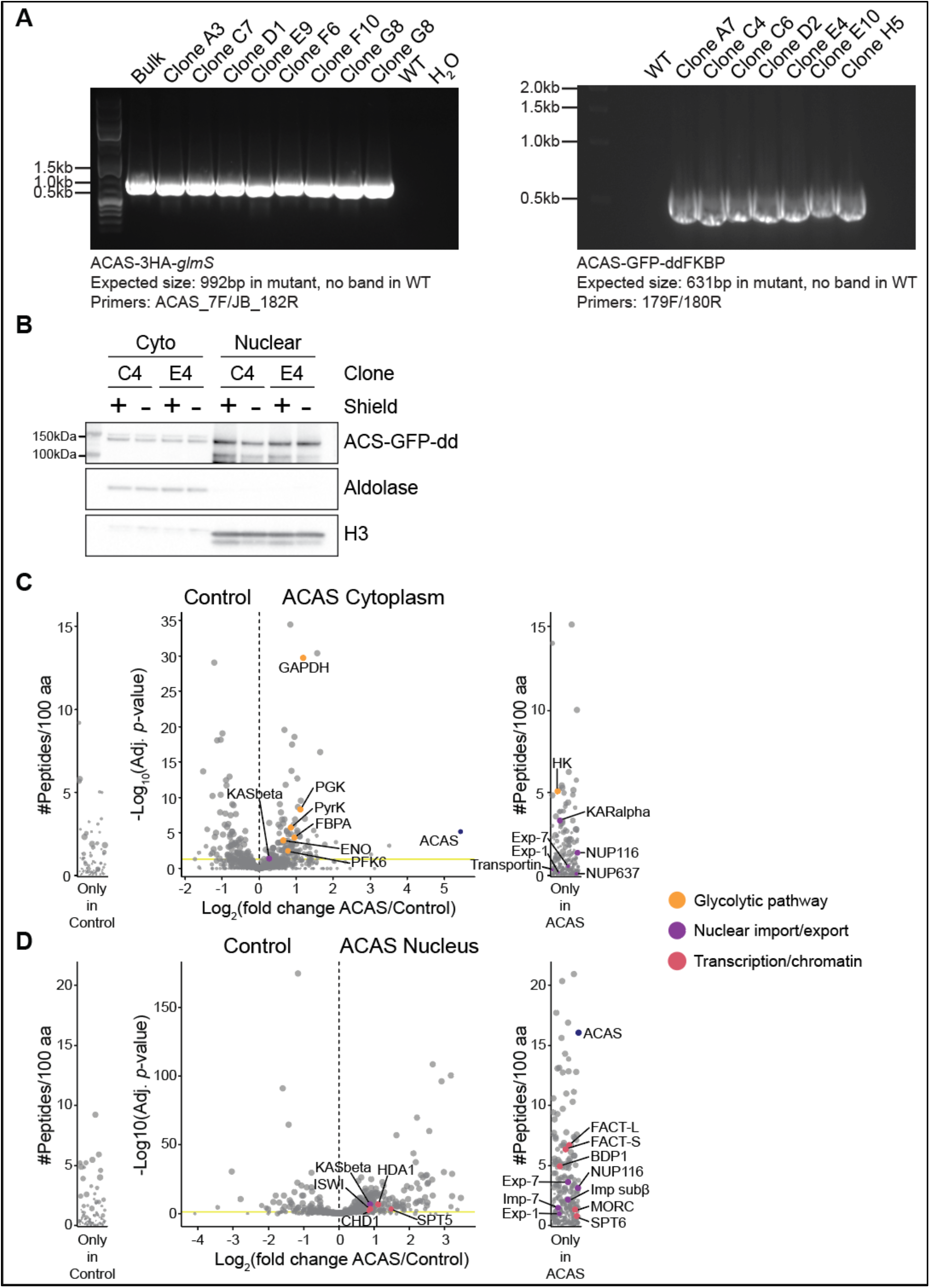
Strain generation and validation. **A.** DNA gels showing PCR validation of the *Pf*ACAS-3HA and *Pf*ACAS-GFP-ddFKBP strains/clones with the indicated primers (**Supp. Data 10**). No genomic DNA (H_2_O) and genomic DNA from WT parasites (WT) were used as controls. DNA size is indicated with a ladder at the left side of each gel and expected band sizes are indicated at the bottom of each gel. **B.** Western blot analysis of nuclear and cytoplasmic (Cyto) fractions from synchronous clonal (C4 and E4) populations of *Pf*ACAS-GFP-ddFKBP (detected with an anti-GFP antibody) parasites in trophozoites [24 hours post invasion (hpi)] in the presence (+) or absence (-) of Shield. Antibodies against aldolase and histone H3 are used as cytoplasmic and nuclear controls, respectively. Molecular weights are shown to the left. **C&D.** *Pf*ACAS IP-MS volcano plot of label-free quantitative proteomic analysis comparing proteins immunoprecipitated with anti-GFP from cytoplasmic (**C**) or nuclear (**D**) fractions in ACAS-GFP parasites compared to wildtype control. Each dot represents a protein, and its size corresponds to the number of peptide ions values used to quantify the ratio of enrichment. x-axis = log_2_(ACAS/Control), y-axis = −log_10_(adjusted *p*-value), vertical dashed line indicates fold change = 1, and horizontal yellow line indicates adjusted *p*-value = 0.05. Side panels show proteins uniquely identified in either sample (y-axis = number of peptides per 100 amino acids). ACAS is indicated in blue, and proteins in the glycolytic pathway, involved in nuclear import/export, and related to transcription and chromatin are indicated in orange, purple, and pink, respectively. All comparisons can be found in **Supp. Data 1 & 2**.

**Expanded View Figure 2:**
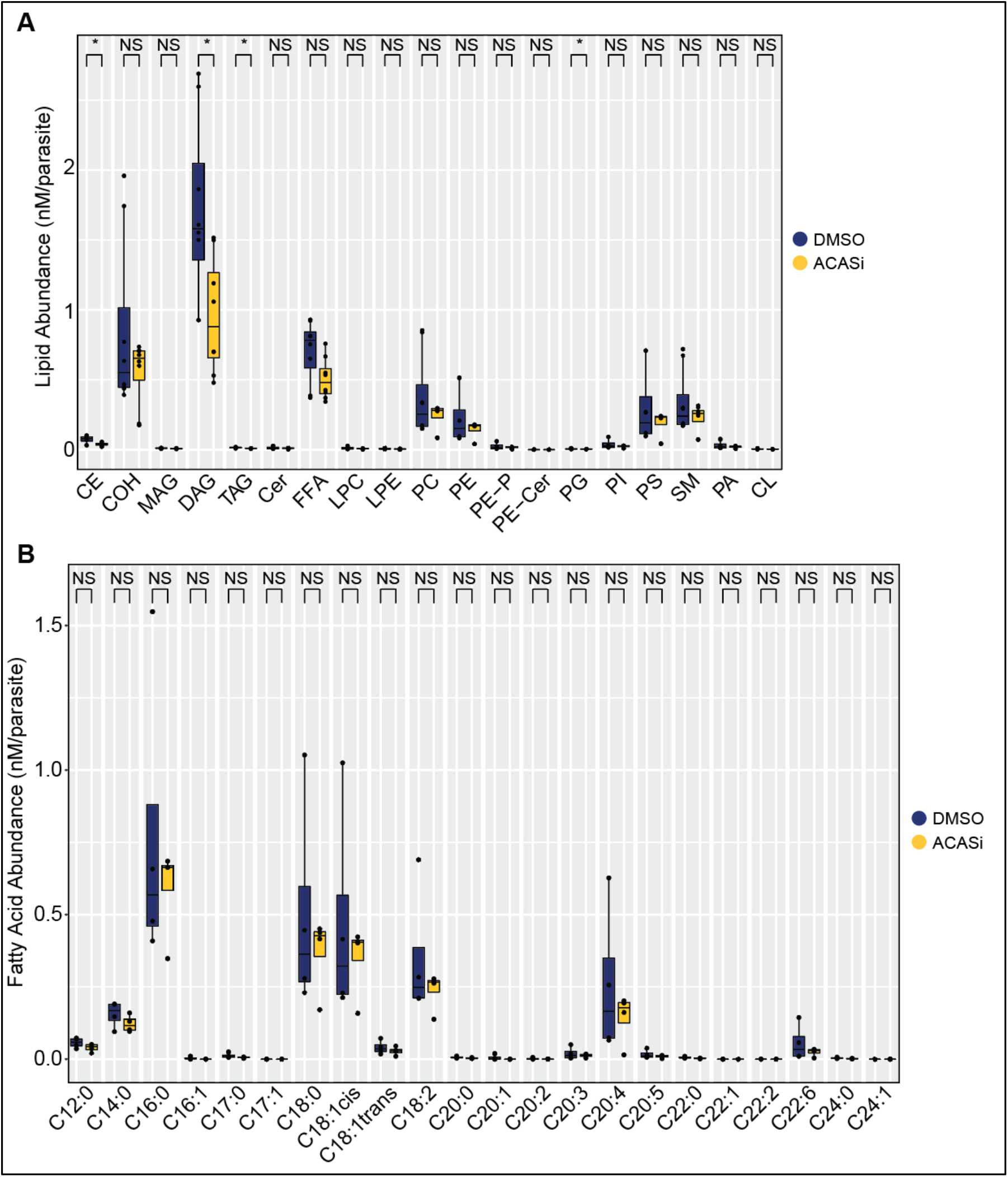
ACAS inhibition induces reduction in some types of lipids. Mass spectrometry (GC-MS) analyses of total lipid class abundance **(A)** and fatty acid abundance **(B)** (nM/parasite), from synchronous wildtype trophozoites treated with DMSO-(blue) or ACASi (yellow) for three hours. Boxes represent the median and interquartile range (IQR), and whiskers represent maximum and minimum values of four replicates. P values are from a Wilcoxon test between ACASi-and DMSO-treated conditions. Lipid abbreviations: CE: cholesteryl-ester, COH: cholesterol, MAG: monoacylglycerol, DAG: diacylglycerol, TAG: triacylglycerol, Cer: ceramide, FFA: free fatty acid, LPC: lyso-phosphatidylcholine, LPE: lyso-phosphatidylethanolamine, PC: phosphatidylcholine, PE: phosphatidylethanolamine, PE-P: phosphatidylethanolamine plasmalogen, PE-Cer: phosphatidylethanolamine ceramide, PG: phosphoglycerol, PI: phosphatidylinositol, PS: phosphatidylserine, SM: sphingomyelin, PA: phosphatidic acid, CL: cardiolipin. Fatty acid nomenclature of CXX:Y (XX: number of carbons, Y: number of insaturations).

